# RNAi-mediated regulation of *alg-3* and *alg-4* coordinates the spermatogenesis developmental program in *C. elegans*

**DOI:** 10.1101/2023.04.06.535930

**Authors:** Cara McCormick, Alicia K. Rogers

**Affiliations:** Department of Biology, University of Texas at Arlington, Arlington, TX, 76019, USA

## Abstract

Coordination of gene regulatory networks is necessary for proper execution of cellular programs throughout development. In *C. elegans,* spermatogenesis and oogenesis occur during different life stages (L4 and adult, respectively) within the hermaphroditic germline tissue. Yet, it remains unclear how these developmental programs are robustly executed, particularly during stressful conditions. Here we show evolutionarily conserved RNA interference (RNAi) pathways act to restrict expression of spermatogenesis genes to the L4 stage. Our findings indicate a novel RNAi-mediated regulatory cascade is essential for properly coordinating the spermatogenesis developmental program, particularly during heat stress. This is achieved through RNAi-mediated genetic switches that regulate the expression of the Argonautes, ALG-3 and ALG-4, to control ALG-3/4 pathway function. Moreover, this work provides key insights into the different molecular mechanisms RNAi pathways employ to maintain both maternal and paternal germ cells’ reproductive potential, and further highlights the complexities and importance of RNAi-mediated gene regulation in development.

## INTRODUCTION

Proper coordination of gene regulatory networks throughout development is required to execute the cellular processes that ensure an animal’s reproductive potential. Environmental or genetic disruptions to gene regulatory networks can have catastrophic effects on physiological processes, such as fertility. In hermaphroditic *Caenorhabditis elegans*, the same germline tissue sequentially completes spermatogenesis and oogenesis during discrete windows of developmental time (the last larval stage (L4) and adult stage, respectively)^1^. While spermatogenesis and oogenesis are temporally separated, sperm and oocytes are derived from the same pools of mitotically replicating nuclei at the distal tip of the gonadal arms, which undergo meiosis as they progress through the germline towards the proximal end of the gonad^2^. Thus, proper coordination of the gene regulatory networks that govern the switch from spermatogenesis to oogenesis within the *C. elegans* germline is essential for maintaining the animal’s maternal and paternal fertility.

Small RNA pathways, also called RNA interference (RNAi) because they often silence target transcripts, are an evolutionarily conserved mechanism that regulates the expression of endogenous and exogeneous genes in all animals. RNAi pathway-mediated gene regulation is critical for maintaining trans-generational epigenetic signals, preserving the germline genome’s integrity from transposons, and safeguarding germ cell identity through maintenance of chromatin states, thus, playing a key role in fertility. Key components of RNAi pathways are RNA-induced silencing complexes (RISC). RISC are made up of an Argonaute protein and its associated small RNA, typically ∼20-30 nucleotides in length, which guides the Argonaute to target transcripts to initiate either transcriptional or post-transcriptional silencing^3–9^. In *C. elegans*, the RNase-III-like enzyme Dicer cleaves exogenous and endogenous long double-stranded RNAs (dsRNAs) to produce ‘primary’ small interfering RNAs (siRNAs), which are 26 nucleotides (nt) in length and have a 5’ guanine (G) (26G-RNAs). These 26G-RNAs form complexes with specific Argonautes to form ‘primary’ RISC that recognize target transcripts^10, 11^. Target recognition by primary RISC then triggers RNA-dependent RNA polymerase (RdRP)-mediated amplification of ‘secondary’ siRNAs from the target transcript. These secondary siRNAs are 22 nt in length with a 5’ G (22G-RNAs) and can be complexed into ‘secondary’ RISC. Amplification of 22G-RNAs is critical for maintaining robust target silencing and can occur in both a *mutator* complex-dependent and independent manner. The *mutator* complex is nucleated around the intrinsically disordered protein, MUT-16, and is housed within perinuclear germ granules called *Mutator* foci, that sit adjacent to the perinuclear hubs for RISC-mediated mRNA surveillance, termed P granules^3, 4, 7, 8, 12^. There are several distinct branches of RNAi pathways (ALG-3/4, CSR-1, ERGO-1, *mutator*, Piwi-interacting RNA (piRNA), and RDE-1). Each branch is defined by the primary and secondary RISC complexes that act on a specific set of targets, the mechanisms of action on targets, and the branch’s dependency on the *mutator* complex for RdRP-mediated amplification of 22G-RNAs. The ERGO-1, *mutator*, piRNA, and RDE-1 RNAi pathway branches rely on MUT-16-dependent 22G-RNA amplification, whereas the ALG-3/4 and CSR-1 branches utilize the RdRP, EGO-1, to achieve MUT-16-independent 22G-RNA amplification^12–14^. The ALG-3/4 and CSR-1 pathways have been shown to be essential for balancing small RNA-mediated positive and negative regulation to permit appropriate expression levels of spermatogenesis-enriched genes during the narrow timeframe of the L4 developmental stage^13, 15–23^.

Mutations in factors of the different RNAi pathway branches can lead to a temperature-sensitive progressive transgenerational loss of fertility, termed the mortal germline phenotype (*mrt*), that manifests at differing generations^3, 12, 15, 24–27^. For example, when animals with a null mutation in the core component of the *mutator* complex, MUT-16, are exposed to heat stress, sterility is trigged within a single generation^14, 24^. When animals are grown at elevated temperature, the totipotent germ cells that give rise to mature sperm and oocytes are constitutively exposed to heat stress during the transition from the L4 stage and adult stage, thus *mrt* phenotypes can be caused by maternal effects, paternal effects, or both within the germ cells. In *C. elegans*, loss of functional MUT-16, and thus amplification of secondary siRNAs, triggers temperature-sensitive sterility caused by both a maternal and paternal effect of *mut-16(pk710)*^24^. The maternal effect of *mut-16* triggers sterility after three generations at elevated temperature (25°C) and is associated with severe morphological gonadal defects, aberrant expression of somatic and spermatogenesis genes due to changes in chromatin accessibility, and ultimately loss of germ cell identity^24^. The paternal effect of *mut-16* results in sterility after a single generation of exposure to heat stress; however, the transcriptional and physiological consequences of the paternal effect of *mut-16* remain unknown. Moreover, it is unclear whether the same or different molecular mechanisms give rise to the heat stress-triggered maternal and paternal consequences within the totipotent germ stem cells from which both oocytes and sperm are both derived.

Here, we demonstrate that MUT-16, and thus *mutator* complex-dependent small RNA amplification, is required to properly coordinate and restrict spermatogenesis gene expression to the L4 developmental stage during heat stress. We leveraged comparative bioinformatic analyses on transcriptional changes in L4 and adult *mut-16* mutant and wild-type animals cultured at permissive (20°C) and elevated (25°C) temperature. Our analyses revealed that in heat stressed *mut-16* mutants, spermatogenesis-enriched genes are down-regulated during the L4 developmental stage, when spermatogenesis typically occurs, and are up-regulated during the adult developmental stage. We found the mis-regulation of these genes, which largely overlapped with ALG-3/4 pathway targets, occurs in a small RNA-dependent manner. These results indicate that MUT-16 is essential for maintaining appropriate small RNA-mediated regulation of spermatogenesis genes throughout development, despite ALG-3/4-class and CSR-1-class 22G-RNAs using MUT-16-independent mechanisms for their amplification. Further analysis revealed that expression of *alg-3* and *alg-4* is developmentally mis-regulated in heat stressed *mut-16* mutants, correlating with changes in 22G-RNA and 26G-RNA levels mapping to the *alg-3* and *alg-4* genomic loci. These findings indicate that small RNA pathways are responsible for switching on and off the ALG-3/4 pathway during the appropriate developmental timeframe and imply that RNAi-mediated cascades may be a conserved gene regulatory network architecture essential for coordinating developmental programs.

In addition to revealing that the *mutator* complex is essential for developmentally regulating *alg-3* and *alg-4*, and thus ALG-3/4 pathway function, we found the physiological manifestation of the paternal *mut-16* effect is severe spermiogenesis defects that result in the onset of sperm-based sterility during heat stress. Moreover, this work reveals that the molecular mechanisms underlying the maternal and paternal effects of *mut-16* during heat stress differs, with the former manifesting via chromatin state modifications (possibly through the nuclear RNAi pathways) and the latter manifesting through loss of small RNA-mediated gene licensing. The identification of sexually dimorphic and temporal differences in regulatory functions of the same RNAi pathway factors within the same gonadal tissue has far-reaching implications for our understanding of regulatory pathways and how they achieve robust, homeostatic gene expression to coordinate cellular programs throughout development.

## RESULTS

### The paternal effect of *mut-16* triggers temperature-sensitive sterility and mild male soma defects

Our previous work established that both maternal and paternal effects of *mut-16(pk710)* lead to temperature-sensitive sterility^24^. To further investigate the paternal effect of *mut-16*, we performed mating brood assays to examine the reproductive potential of *mut-16* males during heat stress. We mated *fog-2(oz40)* hermaphrodites, which are unable to produce their own sperm^28^, and *mut-16* mutant hermaphrodites cultured at 20°C and 25°C, with wild-type (N2) or *mut-16* mutant males cultured at 20°C and 25°C. Males carried the PGL-1::GFP::3xFLAG transgene to ensure only cross progeny were counted. Synchronized L1 animals were cultured at 20°C or 25°C temperature prior to mating plate set up, and crosses were maintained at 25°C if either individual was raised at elevated temperature. Consistent with our previous observations^24^, heat-stressed *mut-16* mutant hermaphrodites, which would be sterile if unmated, produced progeny after mating with wild-type males grown at 20°C and 25°C or *mut-16* mutant males grown at 20°C (Figure 1A). Strikingly, *mut-16* mutant males cultured at 25°C were unsuccessful in producing progeny after mating, regardless of the temperature at which the mother was cultured (Figure 1A). These findings corroborate our previous observation that the paternal effect of *mut-16* manifests after a single generation of elevated temperature^24^, and indicates that sperm-based fertility is more susceptible than oocyte-based fertility to perturbations in small RNA pathways during stressful environmental conditions.

**Figure 1.**
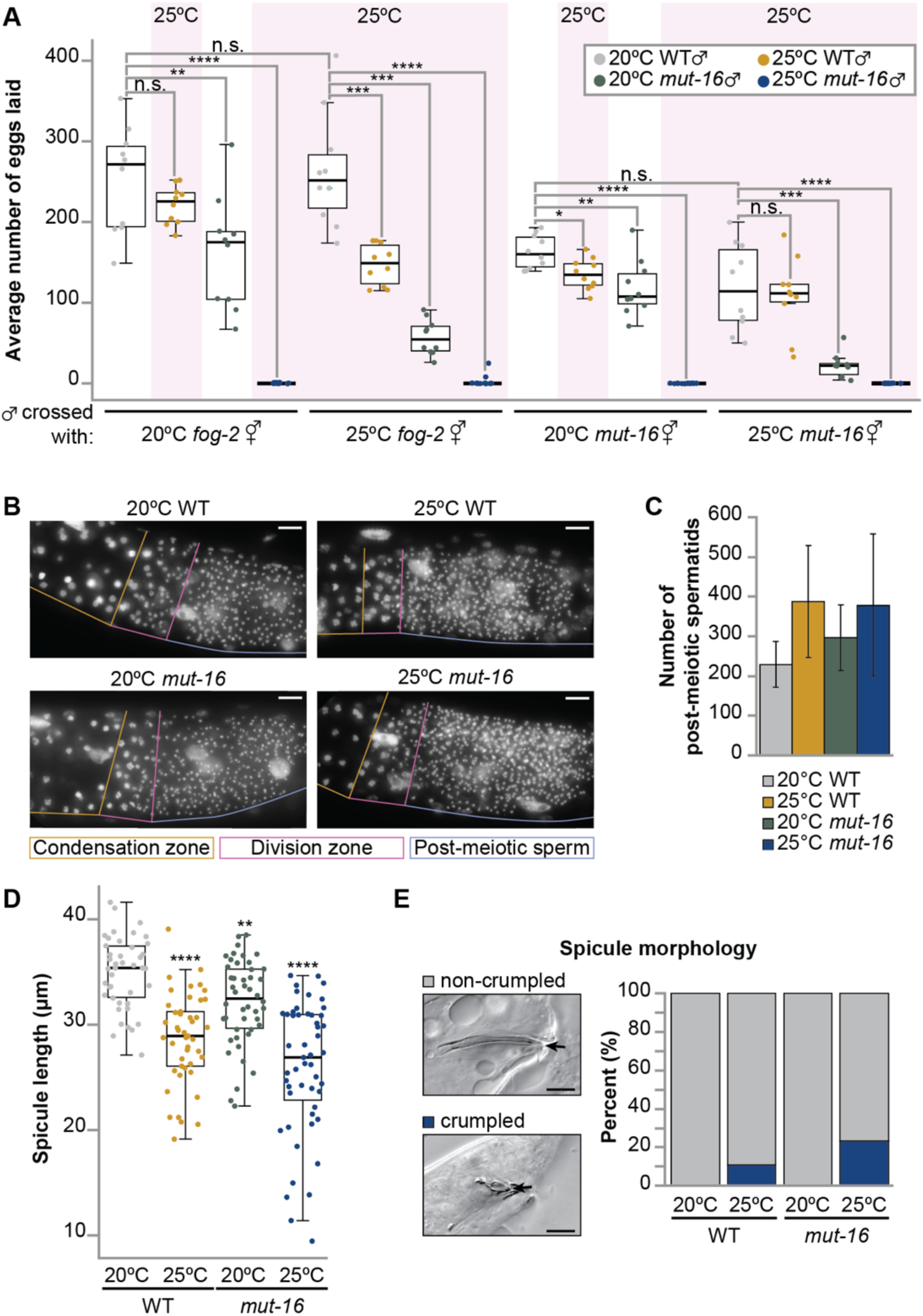
Temperature-sensitive sterility in *mut-16* mutant males correlates with mild somatic defects. **(A)** Shown is average number of eggs laid for *fog-2(oz40*) or *mut-16(pk710)* hermaphrodites cultured at 20°C or 25°C mated with *pgl-1::GFP::3xFLAG* or *mut-16(pk710); pgl-1::GFP::3xFLAG* males cultured at 20°C or 25°C. Circles represent number of GFP-expressing eggs counted for each individual; black line indicates median. n = 10 individuals. Significance compared to mating to wild-type males grown at 20°C is indicated. **(B)** Representative images of DAPI-stained nuclei in male gonads undergoing spermatogenesis of wild-type and *mut-16(pk710)* animals cultured at 20°C or 25°C. Regions of the gonad are labeled: condensation zone (gold), meiotic division zone (pink), and haploid spermatids (blue). Scale bar indicates 10 μm. **(C)** Shown is the average number of post-meiotic haploid spermatids in male gonads of wild-type and *mut-16(pk710)* animals cultured at 20°C or 25°C. Error bars indicate standard deviation. n = 5. **(D)** Shown is spicule length, in μm, measured for wild-type males grown at 20°C (n = 42), wild-type males grown at 25°C (n = 45), *mut-16* mutant males grown at 20°C (n = 46), and *mut-16* mutant males grown at 25°C (n = 51). Circles represent each spicule measured; black line indicates median. Significance compared to wild-type males grown at 20°C is indicated. **(E)** Shown is the percentage of spicules that were normal and non-crumpled (gray) or crumpled (blue) for wild-type males grown at 20°C (n = 42), wild-type males grown at 25°C (n = 45), *mut-16* mutant males grown at 20°C (n = 46), and *mut-16* mutant males grown at 25°C (n = 51). Representative images of non-crumpled and crumpled spicules are shown to the left. Black arrows indicate the tip of the measured spicule. Scale bar indicates 10 μm. n.s. denotes not significant and indicates a p-value > 0.05, * indicates a p-value ≤ 0.05, ** indicates a p-value ≤ 0.01, *** indicates a p-value ≤ 0.001, and **** indicates a p-value ≤ 0.0001.

The maternal effect of *mut-16* during heat stress triggers transgenerational loss of fertility associated with increased germline morphology defects and shrinkage of the gonad^24^. Male infertility can arise from changes in male mating behavior, defects in somatic tissues required for spicule insertion and sperm transfer, or abnormalities in sperm production and/or activation. To identify the cause of the male fertility defect, we first examined the germlines of wild-type or *mut-16* mutant males cultured at 20°C and 25°C for morphological abnormalities and differences in sperm production. We did not observe obvious changes in the germline tissue of *mut-16* mutant males grown at 25°C (Figure 1B). Interestingly, we found that the average number of haploid post-meiotic spermatids in males was increased by the *mut-16* mutation, heat stress, and the combination of both (Figure 1C). The observed increase in spermatid count could point to prolonged spermatogenesis in these animals as a result of a delayed switch from the spermatogenesis to oogenesis developmental program. Next, we assessed the morphology of the somatic spicule tissue, which is critical for probing the vulva and facilitating sperm transfer during mating. Previously, in wild-type male *C. elegans*, it was shown that heat stress results in shortened spicule length and increased incidence of malformed or crumpled spicules^29^. Indeed, we observed increased incidence of malformed/crumpled spicules and overall shorter average spicule lengths in wild-type males grown at 25°C compared to those grown at 20°C (Figure 1D and E). Interestingly, at permissive temperature (20°C), the *mut-16* mutation resulted in an overall shorter average spicule length, but not increased incidence of crumpled spicules; however, *mut-16* mutants that experienced heat stress exhibited further shortening of the average spicule length and increased incidence of malformed/crumpled spicules (Figure 1D and E). Overall, our data indicates that the underlying cause of temperature-sensitive sterility in *mut-16* mutant males is not germline tissue abnormalities or reduced production of spermatids. Heat stress and the *mut-16* mutation do trigger mild defects in the male somatic tissues that facilitate sperm transfer during mating; however, as these defects are not fully penetrant in the population, they are not sufficient to explain the temperature-sensitive sterility of *mut-16* mutant males.

### Down-regulation of spermatogenesis genes correlates with the onset of sperm-based sterility in heat stressed *mut-16* mutants

As germline morphology defects and reduced abundance of spermatids did not underly the onset of temperature-sensitive sterility of *mut-16* mutant males, we next sought to identify molecular changes that dictate the loss of sperm-based fertility. Previously, we found the onset heat stress-induced sterility triggered by the maternal effect of *mut-16* correlates with aberrant expression of spermatogenesis-enriched and somatic genes within the adult germline, when the animal should be undergoing oogenesis^24^. We wondered whether the paternal effect of *mut-16* triggers similar changes in gene expression, so we aimed to identify the mRNA expression changes that occur during heat stress in the L4 developmental stage, when spermatogenesis should occur. To this end, we extracted total RNA from synchronized L4 wild-type and *mut-16* mutants cultured at 20°C and 25°C for a single generation and generated mRNA-seq libraries and size selected small RNA-seq libraries (Table S1). We used differential expression analysis to identify genes whose mRNA levels changed due to elevated temperature (25°C), the *mut-16* mutation, and the combination of both. We identified genes significantly up- or down-regulated (425 and 81 genes, respectively) exclusively in *mut-16* mutants cultured at 25°C. The resulting gene lists omit any genes we identified differentially expressed due to heat stress or the *mut-16* mutation alone. Surprisingly, our bioinformatic analyses revealed that soma-enriched and sperm genes are not up-regulated in heat stressed *mut-16* mutant L4s (Figure 2A). The 425 genes up-regulated exclusively in *mut-16* mutant L4s cultured at 25°C were slightly enriched for oogenesis and gender-neutral germline genes but not somatic (muscle and neuronal) or spermatogenesis genes (Figure 2A and Supplementary Figure S1A). We confirmed muscle and neuronal genes are not up-regulated in L4 *mut-16* mutants using qRT-PCR (Supplementary Figure S1B). These findings suggest aberrant expression of somatic genes within the germline during heat stress is only a consequence of the maternal effect, and not the paternal effect, of *mut-16*. When we performed enrichment analyses on the genes down-regulated exclusively in heat stressed *mut-16* mutant L4s, we found an enrichment of spermatogenesis factors amongst the genes (Figure 2B and C). In addition, tissue enrichment analysis indicated these down-regulated genes are typically expressed in the male tissues of *C. elegans* (Figure 2D). It should be noted that our list of genes exclusively down-regulated upon heat stress in *mut-16* mutants lacks sperm genes whose expression is reduced due to the *mut-16* mutation at 20°C and is further exacerbated at 25°C, 82% of which are members of the major sperm protein (msp) family. Since we were investigating a male fertility defect, the observed dramatic reduction in spermatogenesis gene expression was intriguing. Thus, moving forward in our study of the paternal effect of *mut-16*, we focused on the genes down-regulated exclusively during temperature-sensitive sterility in *mut-16* mutant L4s.

**Figure 2.**
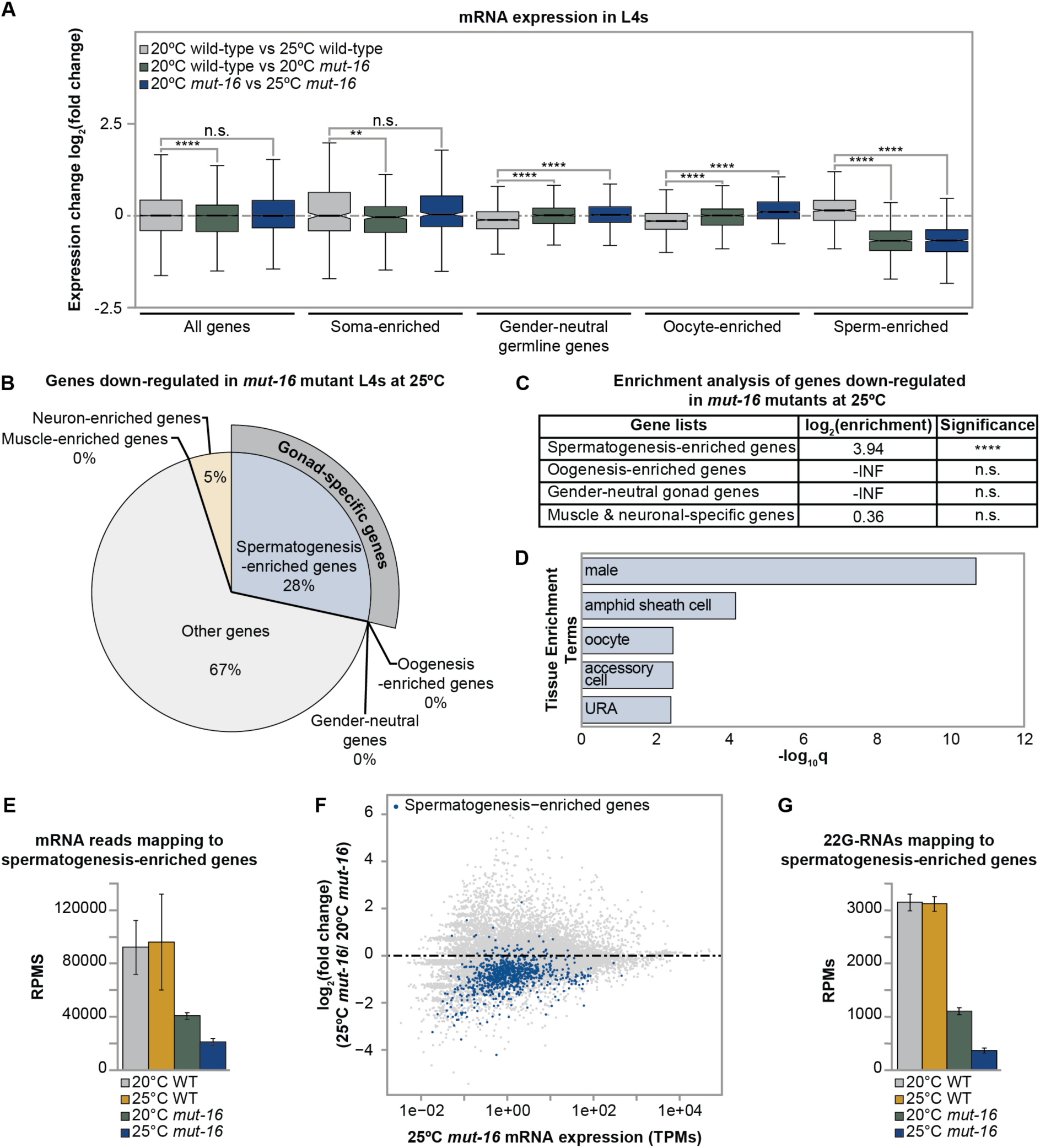
Expression changes for protein-coding genes in heat stressed *mut-16* mutants during the L4 developmental stage. **(A)** Comparison of expression changes in wild-type L4s cultured at 20°C compared to wild-type L4s cultured at 25°C (gray), wild-type L4s cultured at 20°C compared to *mut-16* mutant L4s cultured at 20°C (green), and *mut-16* mutant L4s cultured at 20°C compared to *mut-16* mutant L4s cultured at 25°C (blue) for published enriched gene sets. Notches indicate the 95% confidence interval of the median; black line indicates median. Significance between the log_2_(fold change) upon heat stress in wild-type L4s compared to the log_2_(fold change) due to the *mut-16* mutation and due to the *mut-16* mutation at 25°C is indicated. **(B)** Percentages of gonad-specific and non-gonad-specific genes represented in the genes down-regulated exclusively in *mut-16* mutant L4s at 25°C. **(C)** Enrichment analysis for spermatogenesis, oogenesis, gender-neutral genes, and muscle-specific and neuronal-specific genes amongst the genes down-regulated during heat stress in *mut-16* mutants. **(D)** Tissue enrichment analysis was performed using WormExp (log_10_Q ≥ 2 and FDR < 0.05) for genes down-regulated exclusively in *mut-16* mutants at 25°C. **(E)** mRNA transcripts mapping to spermatogenesis-enriched genes are counted, in reads per million (RPMs), for wild-type and *mut-16* mutant L4 animals cultured at 20°C and 25°C. Error bars indicate standard deviation. **(F)** Shown is difference in expression (log_2_(fold change)) for genes in *mut-16* mutants at 25°C compared to *mut-16* mutants at 20°C plotted according to the average abundance of normalized reads (in transcripts per million (TPMs)) in libraries from *mut-16* mutant L4s grown at 25°C. Each dot represents a gene, with spermatogenic-enriched genes highlighted in blue. **(G)** 22G-RNAs mapping to spermatogenesis-enriched genes are counted, in reads per million (RPMs), for wild-type and *mut-16* mutant L4 animals cultured at 20°C and 25°C. Error bars indicate standard deviation. n.s. denotes not significant and indicates a p-value > 0.05, ** indicates a p-value ≤ 0.01, and **** indicates a p-value ≤ 0.0001.

To further examine the effects on sperm gene expression levels, we assessed the mRNA levels of all spermatogenesis-enriched genes. Previously, we found that heat stress increases expression of sperm genes in wild-type adults^24^. Here, fold change analyses between wild-type L4s grown at 20°C and 25°C indicated that heat stress moderately up-regulates sperm genes (Figure 2A). We corroborated this effect using qRT-PCR with primers for sperm factors that were highly up-regulated in heat stressed wild-type adults (Supplementary Figure S1C). However, it should be noted that assessing the reads per million (RPMs) mapping to all spermatogenesis-enriched genes does not reveal a significant change in RPM levels in wild-type animals at 25°C (Figure 2E). This may indicate that the observed difference in fold change in wild-type animals is driven by a dramatic change in expression of only a subset of sperm genes rather than a global change for all sperm genes. Next, we sought to establish the effects of the *mut-16* mutation on spermatogenesis-enriched gene expression during the L4 developmental stage at 20°C and 25°C. We found that sperm genes are significantly down-regulated in *mut-16* mutants compared to wild-type animals (Figure 2A and E). This contrasts with our previous observation that the *mut-16* mutation leads to up-regulation of sperm genes in the germline of adult animals^24^. We next observed that heat stress exacerbates the reduction in spermatogenesis gene expression (Figure 2A and E-F) and we corroborated this decrease using qRT-PCR with primers for spermatogenesis factors previously observed to be highly up-regulated in adult *mut-16* mutant animals (Supplementary Figure S1D). In adult *C. elegans*, increased expression of spermatogenesis genes is independently triggered by heat stress or loss of MUT-16-dependent 22G-RNAs^24^. Thus, we wanted to determine whether changes in 22G-RNAs mapping to spermatogenesis genes correlated with the observed reduction in their expression in L4 *mut-16* mutants. Our analyses revealed that elevated temperature does not alter 22G-RNA levels mapping to sperm genes in wild-type L4s, but disruption of the *mutator* complex in *mut-16* mutants does trigger reduced 22G-RNA targeting of sperm gene loci that correlates with their down-regulated expression (Figure 2G). Heat stress further exacerbated these effects (Figure 2G). Taken together, our data indicates that during the L4 developmental stage, expression of spermatogenesis genes is susceptible to perturbations in MUT-16-dependent small RNA populations. Furthermore, the growth at elevated temperature worsens the effect of *mut-16*, suggesting that MUT-16-dependent small RNA amplification plays a role in mitigating the effects of heat stress on sperm gene expression.

### Proper expression of spermatogenesis genes throughout development requires MUT-16-dependent small RNAs

In the hermaphroditic germline of *C. elegans*, spermatogenesis and oogenesis occur during specific developmental time points within discrete regions of the same gonadal tissue. As we observed reduced expression of spermatogenesis genes in heat stressed *mut-16* mutant L4s, we wanted to understand whether the paternal effect of *mut-16* triggers mis-coordination of spermatogenesis and oogenesis throughout development at 25°C. To this end, we used our previously published mRNA-seq and small RNA-seq libraries generated from adult wild-type and *mut-16* mutant animals grown at 20°C and 25°C to plot the expression levels of all spermatogenesis-enriched genes and oogenesis-enriched genes across the L4 and adult developmental stages. To account for potential differences when comparing L4 and adult mRNA samples, we normalized the expression levels to *rpl-32*, which encodes a ribosomal protein subunit and is expressed at comparable levels in wild-type and *mut-16* mutant animals throughout the L4 and adult stages (Supplementary Figure S2A and S2B). As expected, we found that, in wild-type animals, spermatogenesis genes are highly expressed during the L4 stage before being down-regulated in the adult animal when oogenesis genes are turned on. Elevated temperature (25°C) did cause a slight increase in both spermatogenesis and oogenesis gene transcript levels during adulthood; however, the trend in expression across developmental stages was not altered (Figure 3A and B). Compared to wild-type animals at 20°C, *mut-16* mutants have reduced expression of sperm genes during the L4 stage and slightly elevated sperm gene expression during the adult stage. Strikingly, heat stress exacerbated the effects of the *mut-16* mutation – causing a further reduction in sperm gene expression during the L4 stage and a dramatic increase in their expression during adulthood (Figure 3A). We did not observe a significant change in the expression levels of oogenesis genes across developmental stages in *mut-16* mutants at 20°C or 25°C (Figure 3B).

**Figure 3.**
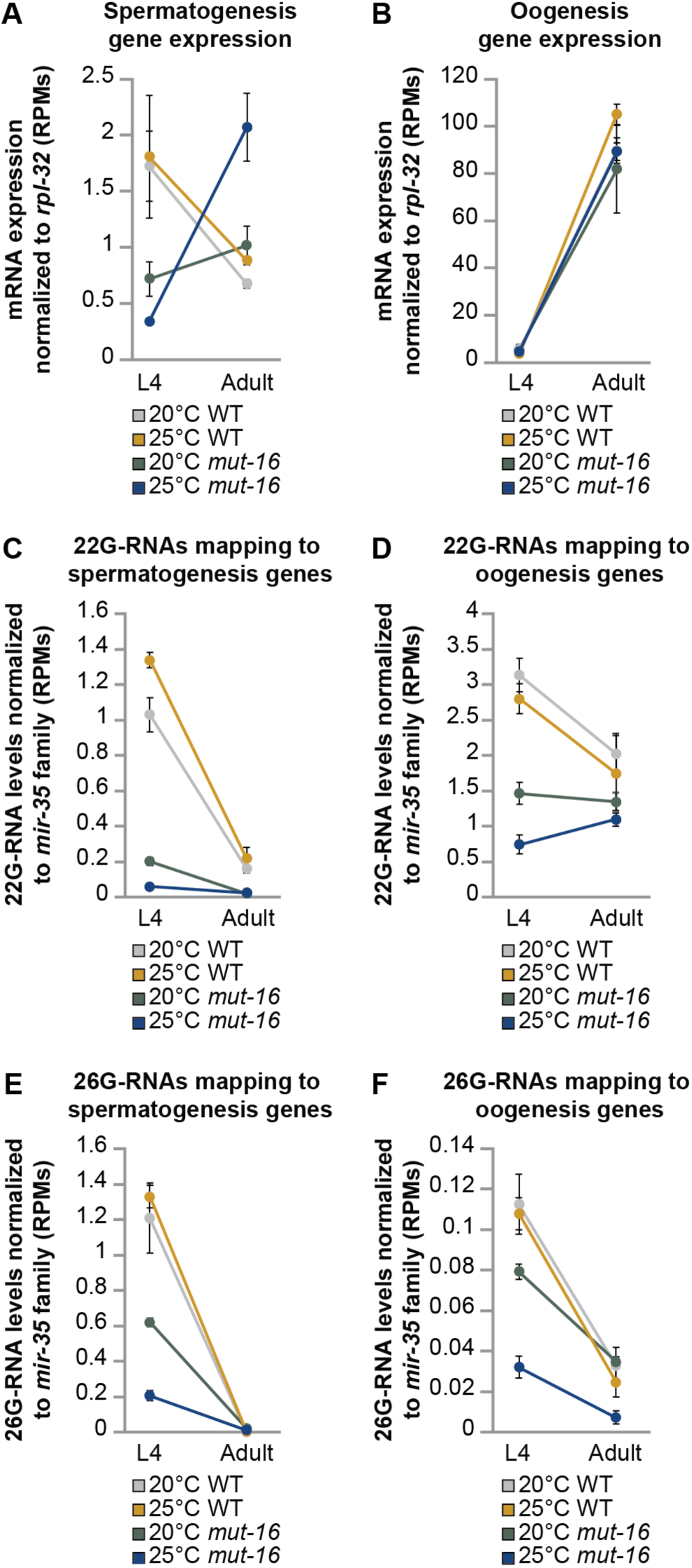
Spermatogenesis genes are developmentally mis-regulated in the absence of MUT-16 during heat stress. **(A)** mRNA transcripts mapping to spermatogenesis-enriched genes are counted and normalized to the expression level of *rpl-32*, in reads per million (RPMs), for L4 and adult wild-type and *mut-16* mutant animals cultured at 20°C and 25°C. Error bars indicate standard deviation. **(B)** mRNA transcripts mapping to oogenesis-enriched genes are counted and normalized to the expression level of *rpl-32*, in reads per million (RPMs), for L4 and adult wild-type and *mut-16* mutant animals cultured at 20°C and 25°C. Error bars indicate standard deviation. **(C)** 22G-RNAs mapping to spermatogenesis-enriched genes are counted and normalized to reads mapping to the *mir-35* family, in reads per million (RPMs), for L4 and adult wild-type and *mut-16* mutant animals cultured at 20°C and 25°C. Error bars indicate standard deviation. **(D)** 22G-RNAs mapping to oogenesis-enriched genes are counted and normalized to reads mapping to the *mir-35* family, in reads per million (RPMs), for L4 and adult wild-type and *mut-16* mutant animals cultured at 20°C and 25°C. Error bars indicate standard deviation. **(E)** 26G-RNAs mapping to spermatogenesis-enriched genes are counted and normalized to reads mapping to the *mir-35* family, in reads per million (RPMs), for L4 and adult wild-type and *mut-16* mutant animals cultured at 20°C and 25°C. Error bars indicate standard deviation. **(F)** 26G-RNAs mapping to oogenesis-enriched genes are counted and normalized to reads mapping to the *mir-35* family, in reads per million (RPMs), for L4 and adult wild-type and *mut-16* mutant animals cultured at 20°C and 25°C. Error bars indicate standard deviation.

Next, we assessed 22G-RNA and 26G-RNA populations mapping to spermatogenesis-enriched and oogenesis-enriched genes to determine whether changes in the small RNA levels correlated with the observed changes in mRNAs. We normalized the levels of 22G-RNAs and 26G-RNAs to all small RNA reads mapping to the germline-specific *mir-35* family (*mir-35-42*), which are not amplified in the *mutator* complex. Our analyses revealed that 22G-RNAs mapping to spermatogenesis and oogenesis genes are strongly depleted throughout development in *mut-16* mutants and that this effect is exacerbated by heat stress (Figure 3C and D). This result is expected as *mut-16* mutants lack the ability to perform *mutator* complex-dependent 22G-RNA amplification; however, it is noteworthy that heat stress worsens the loss of 22G-RNAs. This would suggest that the *mutator* complex plays an important role in mitigating the effects of heat stress on the pools of 22G-RNAs synthesized by *mut-16*-independent mechanisms. Interestingly, 26G-RNAs mapping to both spermatogenesis and oogenesis genes in *mut-16* mutants were reduced during the L4 stage. Heat stress caused further loss of 26G-RNAs mapping to both gene groups during the L4 stage and a reduction in 26G-RNAs targeting oogenesis genes during adulthood (Figure 3E and F). As the *mutator* complex functions downstream of 26G-RNAs, it is puzzling that their levels are impacted in *mut-16* mutants (see discussion). Ultimately, our analyses indicate that the expression profile of oogenesis-enriched genes throughout development is not significantly impacted by elevated temperature (25°C), changes in small RNA populations in *mut-16* mutants, or the combination of both. In contrast, our analyses indicate that the effect of the *mut-16* mutation on spermatogenesis-enriched gene expression is dictated by the developmental stage of the animal. MUT-16, and thus the *mutator* complex amplification of 22G-RNAs promotes expression of sperm genes during the L4 stage and then inhibits their expression during adulthood. This is the first evidence implicating MUT-16-dependent 22G-RNAs in promoting expression of spermatogenesis genes. Moreover, our data suggests that despite the spermatogenesis to oogenesis switch occurring within the same gonadal tissue, these developmental programs are independently regulated at the molecular level.

### Spermiogenesis defects correlate with heat stress-induced sterility in *mut-16* mutants

Loss of sperm’s reproductive potential can occur due to perturbations during spermatogenesis, spermiogenesis, or at the fertilization stage. We next wanted to know whether we could identify stage(s) at which sperm development, and thus sperm-based fertility, might be negatively impacted by differential gene expression in heat stressed *mut-16* mutant L4s. To this end, we assessed the changes in expression of genes critical for proper completion of I) primary spermatocyte division to form secondary spermatocytes, II) spermatid formation, III) spermatid activation during spermiogenesis, IV) spermatozoa entering the oocyte during fertilization, and V) post-fertilization viability of the embryo (Figure 4A). During spermatogenesis, a primary spermatocyte, derived from totipotent germ cells, undergoes rounds of meiotic division to form sessile, round spermatids^30^. Abundant post-meiotic spermatids were present in the germlines of *mut-16* males grown at 25°C (Figure 1B and C), suggesting defects in spermatocyte and spermatid formation do not the underly these animals’ loss of sperm-based fertility. In agreement with this, we found genes critical for proper division of spermatocytes and spermatid formation are mildly, but not statistically significantly (p-value ≤ 0.05), down-regulated in heat stressed *mut-16* mutants (Figure 4B). Spermatids transform into motile spermatozoa through the process of spermiogenesis, during which exposure to an activator causes a spermatid to extend long spikes that are restructured to form a pseudopod^30, 31^. Our differential expression analysis revealed that all the genes essential for proper spermiogenesis had reduced expression in heat stressed *mut-16* mutants, but only *fer-1* and *swm-1* were significantly down-regulated (Figure 4B). It should be noted that *swm-1* was also significantly down-regulated at 25°C in wild-type animals, indicating its expression is particularly susceptible to heat stress (Figure 4B). Loss of functional FER-1 triggers formation of short, nonmotile pseudopods during spermiogenesis^32–35^, whereas spermatids in *swm-1* mutants prematurely activate prior to mating^36^. Mutations in the *spe-8* pathway (*spe-8*, *spe-12*, *spe-27*, and *spe-29*), which is critical for *in vivo* and *in vitro* spermatid activation, result in spermatozoa arresting as nonmotile, spiky intermediates^15, 30, 31^. All of these spermiogenesis defects leads to sperm-based infertility. After spermiogenesis, motile spermatozoa must be able to recognize and adhere/fuse with oocytes to complete fertilization. Our analysis of the expression levels of genes required for fertilization and post-fertilization viability of the embryo found that only *spe-11* was significantly differentially expressed (Figure 4B). Spermatozoa lacking *spe-11* can fertilize oocytes to produce embryos; however, after these eggs are laid, embryogenesis fails, resulting in paternal-effect embryonic lethality^30^. In our mating brood size assay, *fog-2* hermaphrodites crossed to heat stressed *mut-16* mutants males did not lay eggs (Figure 1A). The absence of egg laying upon mating suggests that fertilization is not occurring, and thus *spe-11*-induced paternal-effect embryonic lethality is not likely to be the driving force of temperature-sensitive sperm-based sterility in *mut-16* males.

**Figure 4.**
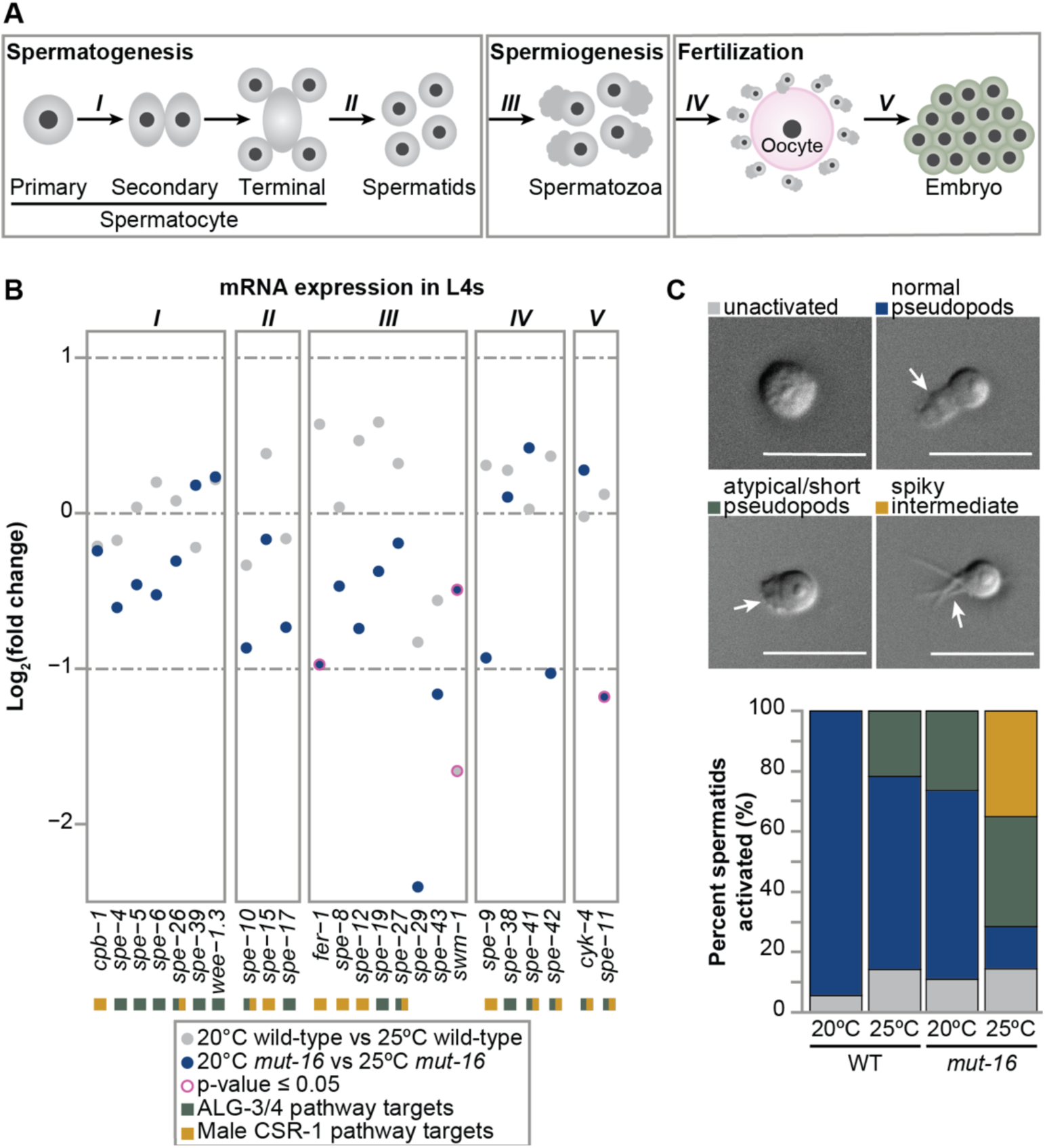
The paternal effect of *mut-16* triggers spermiogenesis defects. **(A)** Schema of stages of sperm development critical for male reproductive potential. I) primary spermatocyte division to form secondary spermatocytes, II) spermatid formation, III) spermatid activation during spermiogenesis, IV) spermatozoa enter the oocyte during fertilization, and V) post-fertilization viability of the embryo **(B)** Strip plot showing the change in expression of genes essential for each stage of sperm development in wild-type L4s cultured at 20°C compared to wild-type L4s cultured at 25°C (gray) and *mut-16* mutant L4s cultured at 20°C compared to *mut-16* mutant L4s cultured at 25°C (blue). Pink ring around data point indicates expression changes with a significant p-value as determined by DESeq2 (p-value ≤ 0.05). Boxes next to gene names indicate whether the gene is targeted by the ALG-3/4 pathway (green) and/or the male CSR-1 pathway (gold). **(C)** Shown are representative images of spermatids after exposure to pronase E. Below, percent of spermatids that are unactivated (gray), have normal pseudopod formation (blue), atypical pseudopod formation (green), or arrest as spiky intermediates (gold) are shown for wild-type and *mut-16* males grown at 20°C and 25°C. White arrows indicate the pseudopods and scale bars indicate 10 μm. n = 200 per genotype for each condition.

Our bioinformatics analyses hinted at spermiogenesis defects underlying heat stress-induced infertility triggered by the paternal effect of *mut-16*. To test this, we performed an *in vitro* activation assay on spermatids obtained from wild-type and *mut-16* mutant males grown to adulthood at 20°C and 25°C and assessed pseudopod formation after exposure to the activation agent, pronase E. We classified pseudopod formation as normal, atypical (short or two distinct pseudopods), or as spiky intermediates (Figure 4C). At permissive temperature (20°C), wild-type spermatids had a 95% activation rate, with all pseudopods forming normally. Heat stress lowered the activation rate of wild-type spermatids to 86% and increased the incidence of atypical pseudopod formation (22% short pseudopods) (Figure 4C and Table S3). Spermatids collected from *mut-16* mutant males at 20°C exhibited similar rates of activation (89%) and incidence of atypical pseudopod formation (27%) to heat stressed wild-type spermatids (Figure 4C and Supplementary Table 3). Intriguingly, spermatids collected from *mut-16* males grown at 25°C had an 86% activation rate, but these largely formed atypical pseudopods or arrested as spiky intermediates (37% and 35%, respectively) (Figure 4C and Table S3). These data indicates that heat stress and the *mut-16* mutation independently trigger comparable spermiogenesis defects, but the combination of both leads to severely compromised pseudopod formation. Taken together, our data indicates that spermiogenesis defects are the physiological manifestation of *mut-*16’s paternal effect. Our mRNA-seq analyses and the specific pseudopod phenotypes observed point to reduced expression of *fer-1* and deficiencies in the expression levels of *spe-8* pathway factors causing the myriad of spermiogenesis defects underlying loss of sperm-based fertility during heat stress in *mut-16* mutants.

### Targets of the *mutator* complex-independent ALG-3/4 pathway are down-regulated in *mut-16* mutants during heat stress

Several distinct small RNA pathway branches (piRNA, ALG-3/4, and CSR-1) have been previously shown to target sperm genes; however, of these, only the piRNA pathway uses the *mutator* complex for 22G-RNA amplification^12–15, 17, 23, 27, 37^. Thus, we hypothesized the spermatogenesis genes down-regulated in a small RNA-dependent manner in heat stressed *mut-16* mutants would be piRNA pathway targets. To test this, we assessed the enrichment of the different RNAi pathway branches’ targets amongst the 81 genes down-regulated exclusively in *mut-16* mutants at 25°C. Surprisingly, these genes were not enriched for targets of the piRNA pathway, but rather for targets of the *mutator* complex-independent ALG-3/4 and male CSR-1 pathways (Figure 5A and B). Because the *mutator* complex canonically does not function in the ALG-3/4 or CSR-1 pathways we sought to further characterize the impacts of heat stress, loss of the *mutator* complex, and the combination of both on these pathways’ targets.

**Figure 5.**
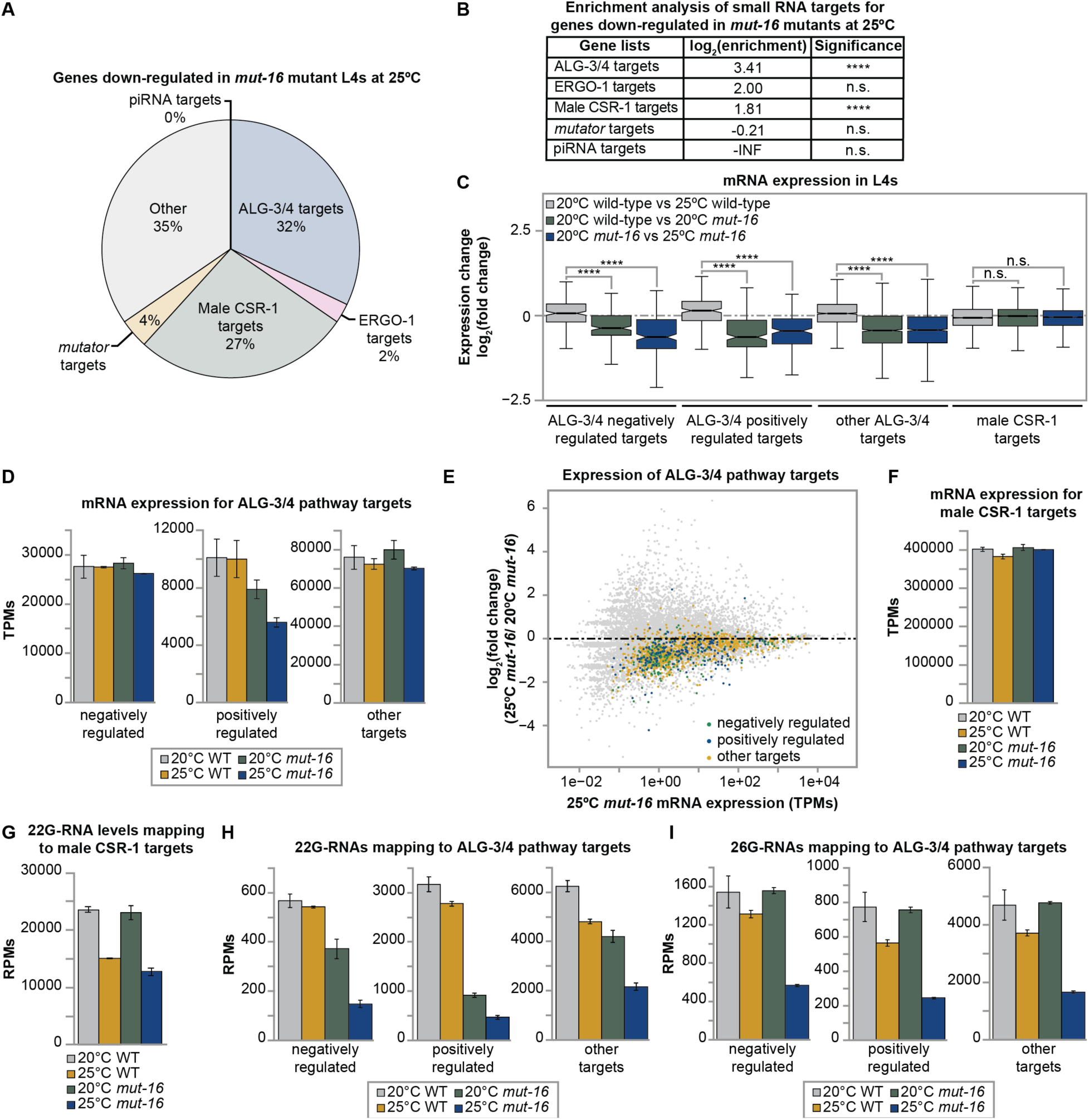
Heat-stressed *mut-16* mutant L4 animals exhibit reduced ALG-3/4 pathway target expression. **(A)** Percentages of genes targeted by distinct small RNA pathways (ALG-3/4, ERGO-1, male CSR-1, *mutator*, and piRNA pathways) represented in the genes down-regulated exclusively in *mut-16* mutant L4s at 25°C. **(B)** Enrichment analysis for ALG-3/4, ERGO-1, male CSR-1, *mutator*, and piRNA pathway targets amongst the genes down-regulated during heat stress in *mut-16* mutants. **(C)** Comparison of expression changes in wild-type L4s cultured at 20°C compared to wild-type L4s cultured at 25°C (gray), wild-type L4s cultured at 20°C compared to *mut-16* mutant L4s cultured at 20°C (green), and *mut-16* mutant L4s cultured at 20°C compared to *mut-16* mutant L4s cultured at 25°C (blue) for published ALG-3/4 pathway and male CSR-1 pathway target gene lists. Notches indicate the 95% confidence interval of the median; black line indicates median. Significance between the log_2_(fold change) at upon heat stress in wild-type L4s compared to the log_2_(fold change) due to the *mut-16* mutation and due to the *mut-16* mutation at 25°C is indicated. **(D)** mRNA transcripts mapping to negatively regulated, positively regulated, and other ALG-3/4 pathway target genes are counted, in transcripts per million (TPMs), for wild-type and *mut-16* mutant L4 animals cultured at 20°C and 25°C. Error bars indicate standard deviation. **(E)** Shown is difference in expression (log_2_(fold change)) for genes in *mut-16* mutants at 25°C compared to *mut-16* mutants at 20°C plotted according to the average abundance of normalized reads (in transcripts per million (TPMs)) in libraries from *mut-16* mutant L4s grown at 25°C. Each dot represents a gene, with negatively regulated ALG-3/4 pathway targets (green), positively regulated ALG-3/4 pathway targets (blue), and other ALG-3/4 pathway targets (gold) highlighted. **(F)** mRNA transcripts mapping to male CSR-1 pathway target genes are counted, in transcripts per million (TPMs), for wild-type and *mut-16* mutant L4 animals cultured at 20°C and 25°C. Error bars indicate standard deviation. **(G)** 22G-RNAs mapping to male CSR-1 pathway target genes are counted, in reads per million (RPMs), for wild-type and *mut-16* mutant L4 animals cultured at 20°C and 25°C. Error bars indicate standard deviation. **(H)** 22G-RNAs mapping to negatively regulated, positively regulated, and other ALG-3/4 pathway target genes are counted, in reads per million (RPMs), for wild-type and *mut-16* mutant L4 animals cultured at 20°C and 25°C. Error bars indicate standard deviation. **(I)** 26G-RNAs mapping to negatively regulated, positively regulated, and other ALG-3/4 pathway target genes are counted, in reads per million (RPMs), for wild-type and *mut-16* mutant L4 animals cultured at 20°C and 25°C. Error bars indicate standard deviation. n.s. denotes not significant and indicates a p-value > 0.05, and **** indicates a p-value ≤ 0.0001.

ALG-3/4 pathway targets can been classified by the effect of ALG-3/4-targeting on their expression: some are negatively regulated by ALG-3/4 while others are positively regulated, and some targets can be immunoprecipitated with ALG-3/4 but do not have altered expression upon loss of the Argonautes^16^. We performed differential expression analysis to assess the transcript levels of all genes categorized as ALG-3/4 negatively regulated targets, ALG-3/4 positively regulated targets, ‘other’ ALG-3/4 pathway targets, and male CSR-1 pathway targets (204, 214, 988, and 2801 genes, respectively). We found that heat stress results in an overall slight up-regulation of all three groups of ALG-3/4 targets. However, it should be noted that a significant change in expression upon heat stress is not observed when assessing TPMs mapping to each ALG-3/4 target group, thus the observed fold change may be driven by only a subset of genes rather than a global change for these target groups (Figure 5C and D). In contrast, the *mut-16* mutation results in overall down-regulation for the three groups of ALG-3/4 targets, with positively regulated targets being the only group to also show a significant reduction in TPM levels (Figure 5C and D). Elevated temperature (25°C) further exacerbates the effect of the *mut-16* mutation on all three ALG-3/4 pathway target groups (Figure 5C-E). ALG-3/4 pathway targets were significantly depleted from the list of genes exclusively up-regulated in *mut-16* mutant L4s at 25°C, further corroborating the negative effect of the *mut-16* mutation on their expression (Supplementary Figure S3A). Expression of male CSR-1 pathway targets was slightly down-regulated upon heat stress in wild-type animals but was not affected by the *mut-16* mutation or the combination of both (Figure 5C and F). This was surprising as male CSR-1 pathway targets were enriched amongst the genes down-regulated exclusively in *mut-16* mutants at 25°C; however, ALG-3/4 pathway targets and male CSR-1 pathway targets overlap considerably (93 male CSR-1 and ALG-3/4 negatively regulated targets, 124 male CSR-1 and ALG-3/4 positively regulated targets, and 581 male CSR-1 and other ALG-3/4 pathway targets). To ensure any changes in gene expression were not masked by the overlapping nature of these two RNAi pathways, we performed the differential expression analysis using gene lists with and without overlapping ALG-3/4 and male CSR-1 pathway targets. However, using non-overlapping gene lists did not change the overall effects observed during our analyses (Supplementary Figure S3B-D). We looked more closely at the male CSR-1 pathway targets present in the list of genes down-regulated specifically in *mut-16* mutants at 25°C and found that most of them (17 out of 22 genes) are also annotated as ALG-3/4 pathway targets. Thus, the overlapping annotations of these genes explains why we find male CSR-1 targets enriched amongst down-regulated genes but do not see overall expression changes for male CSR-1 targets. Ultimately, our differential expression analyses indicate that MUT-16 plays a role in maintaining appropriate expression of positively regulated ALG-3/4 targets at permissive temperature (20°C), and that all three groups of ALG-3/4 targets are sensitive to loss of the *mutator* complex during heat stress.

Next, we sought to determine how changes in mRNA levels correlated with small RNAs mapping to targets of the ALG-3/4 and male CSR-1 pathways. We first assessed 22G-RNA levels mapping to target loci as the *mutator* complex functions to amplify secondary siRNAs. As expected, 22G-RNA levels mapping to the *mutator* complex-independent CSR-1 pathway targets were not impacted by loss of *mut-16* (Figure 5G). However, heat stress led to comparable reductions of 22G-RNAs mapping to male CSR-1 targets in wild-type and *mut-16* mutant animals (Figure 5G). This indicates CSR-1-loaded 22G-RNAs are susceptible to elevated temperature. When we looked at 22G-RNAs mapping to the three groups of ALG-3/4 targets, we found that heat stress triggers a slight reduction in their levels, whereas the *mut-16* mutation caused moderate to severe reductions in 22G-RNA levels mapping to all three groups of targets, with positively regulated targets exhibiting the most severe loss (Figure 5H). The combination of heat stress and loss of *mutator* complex-dependent 22G-RNA amplification had an additive effect, resulting in severely reduced numbers of 22G-RNAs mapping to all three groups of ALG-3/4 targets (Figure 5H). Together, these analyses indicate that 22G-RNAs targeting male CSR-1 and ALG-3/4 pathway targets are susceptible to heat stress. Furthermore, this is the first evidence indicating that a portion of the 22G-RNAs mapping to ALG-3/4 targets, most notably positively regulated ALG-3/4 targets, are amplified in a *mutator* complex-dependent manner. Canonically, 22G-RNAs are generated from ALG-3/4 pathway target mRNAs in a *mutator* complex-independent manner after recognition by ALG-3/4 complexed with a 26G-RNA. To further understand how fluctuations in small RNA populations drive changes in expression of ALG-3/4 pathway targets that correlate with the temperature sensitive sterility in *mut-16* mutants, we then examined levels of the upstream 26G-RNAs mapping to the three target groups. We found that heat stress, but not the *mut-16* mutation alone, caused slight loss of 26G-RNAs mapping to the three groups of ALG-3/4 targets (Figure 5I). However, the combination of heat stress and the *mut-16* mutation resulted in a severe loss of 26G-RNAs mapping to all three ALG-3/4 target groups (Figure 5I). The additive effect of heat stress and loss of *mut-16* on ALG-3/4 pathway 26G-RNA levels suggests the *mutator* complex plays a critical role in mitigating the consequences of elevated temperature on ALG-3/4 pathway function. Taken together, our small RNA-seq analyses indicate that the *mutator* complex (likely indirectly) contributes to maintaining homeostatic levels of 22G-RNAs and 26G-RNAs mapping to ALG-3/4 pathway targets during the L4 developmental stage during stressful conditions.

### Developmental stage-specific expression of *alg-3* and *alg-4* is maintained by *mutator* complex-dependent small RNAs during heat stress

26G-RNA-mediated recognition of target transcripts occurs upstream of 22G-RNA amplification, thus the loss of 26G-RNAs mapping to ALG-3/4 pathway targets in *mut-16* mutants is unexpected. However, we previously showed that loss of *mut-16* triggers a reduction in ERGO-1-class 26G-RNAs through a feedback loop that fine-tunes expression of a factor essential for the biogenesis of ERGO-1-class 26G-RNAs^38^. To further understand the mechanism underlying loss of small RNAs mapping to ALG-3/4 pathway targets in heat stressed *mut-16* mutants, we assessed the expression of known ALG-3/4 pathway factors. We found the two primary Argonaute proteins, ALG-3 and ALG-4, had significantly reduced expression in *mut-16* mutants experiencing heat stress (Figure 6A and B, and Supplementary Figure S4A and B). Using RT-qPCR, we confirmed the observed loss of *alg-3* and *alg-4* expression upon heat stress in *mut-16* mutant L4s (Figure 6C). Next, we examined the small RNA reads mapping to *alg-3* and *alg-4* to determine if these genes are targeted by small RNAs, and whether changes in their expression correlated with differential small RNA targeting. We found that in wild-type animals, small RNAs target both *alg-3* and *alg-4* (Figure 6D and Supplementary Figure S4A and B). Heat stress and the *mut-16* mutation each independently triggered slight reductions in small RNA levels mapping to *alg-3* and *alg-4*; however, the combination of both resulted in a severe loss of small RNAs mapping to these genomic loci (Figure 6D and Supplementary Figure S4A and B). To further characterize the small RNAs targeting *alg-3* and *alg-4*, we examined their read lengths and first nucleotide (nt) biases. We found that these loci are predominately targeted by a group of 22G-RNAs and a group of 26G-RNAs (Figure 6E-G). It should be noted that 21-nt reads mapping to *alg-3* and *alg-4* largely have a 5’ G bias, suggesting these are 22G-RNAs that may have been shortened to 21-nt during reading trimming (Figure 6F and G, and Supplementary Figure S4C). In wild-type animals, elevated temperature (25°C) resulted in a strong loss of 22G-RNAs, but not 26G-RNAs, targeting *alg-3* and *alg-4* (Figure 6F and G, and Supplementary Figure S4A and B). The *mut-16* mutation alone triggered a minor loss of 22G-RNAs and 26G-RNAs targeting *alg-3* and *alg-4*; however, the combination of heat stress and the *mut-16* mutation resulted in severe depletion of both 22G-RNAs and 26G-RNAs mapping to *alg-3* and *alg-4* (Figure 6F and G, and Supplementary Figure S4A and B). This data indicates that L4 stage expression of *alg-3* and *alg-4* is regulated in a small RNA-dependent manner. Moreover, our data suggests that the *mutator* complex is critical for maintaining small RNA-mediated regulation of *alg-3* and *alg-4* during stressful conditions.

**Figure 6.**
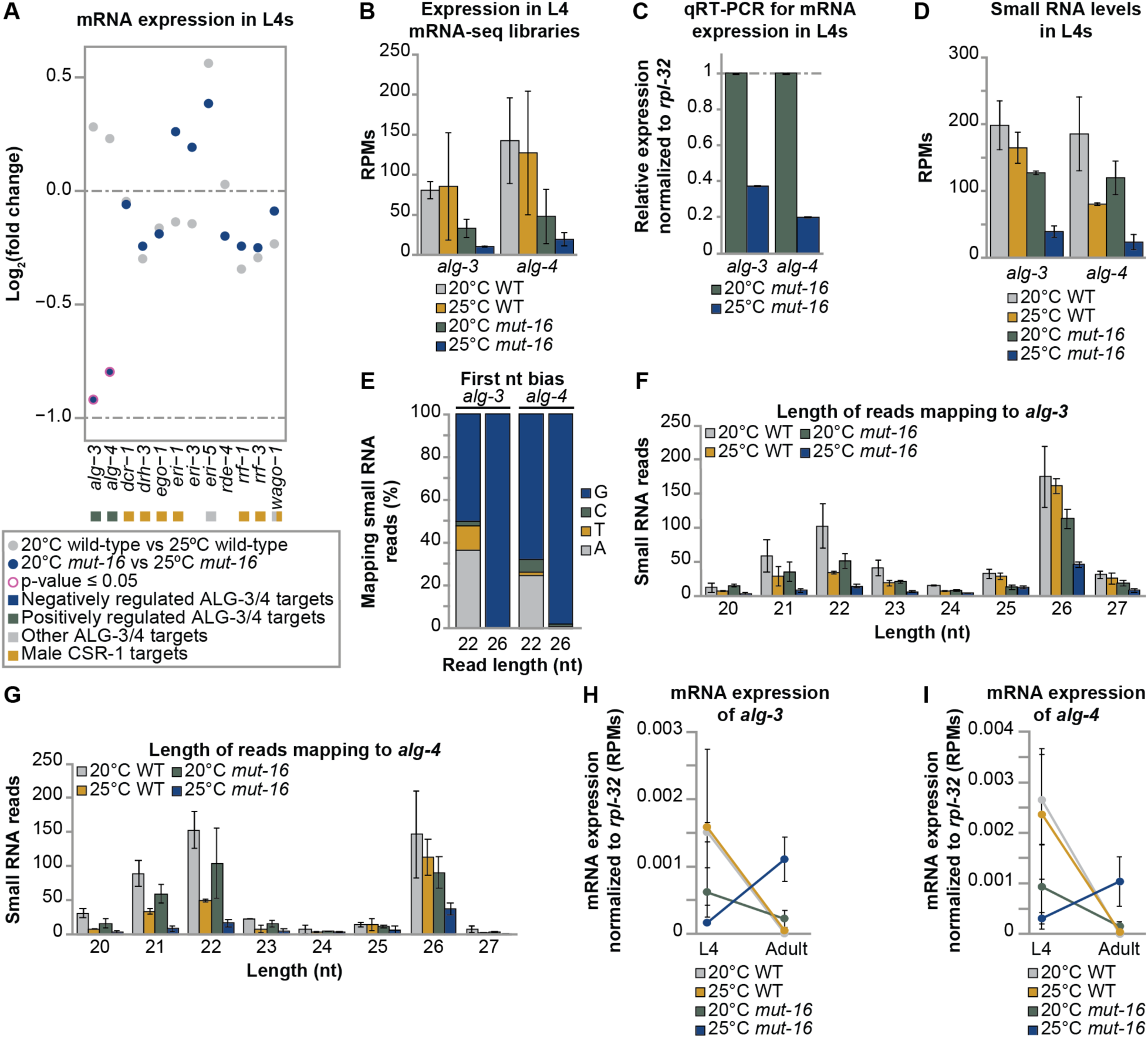
The *mutator* complex is required for developmentally coordinating *alg-3* and *alg-4* expression during heat stress. **(A)** Strip plot showing the change in expression of ALG-3/4 pathway factors in wild-type L4s cultured at 20°C compared to wild-type L4s cultured at 25°C (gray) and *mut-16* mutant L4s cultured at 20°C compared to *mut-16* mutant L4s cultured at 25°C (blue). Pink ring around data point indicates expression changes with a significant p-value as determined by DESeq2 (p-value ≤ 0.05). Boxes next to gene names indicate whether the gene is targeted by the ALG-3/4 pathway (targets negatively regulated in blue, targets positively regulated in green, and other targets in gray) and/or the male CSR-1 pathway (gold). **(B)** mRNA transcripts mapping to *alg-3* and *alg-4* are counted, in reads per million (RPMs), for wild-type and *mut-16* mutant L4 animals cultured at 20°C and 25°C. Error bars indicate standard deviation. **(C)** qRT-PCR assay of *alg-3* and *alg-4* expression, normalized to *rpl-32* expression, in *mut-16* mutant L4 animals cultured at 25°C normalized to expression levels in *mut-16* mutant L4 animals cultured at 20°C. Error bars indicate standard deviation. n = 3 biological replicates. **(D)** Total small RNA reads per million (RPMs) mapping to *alg-3* and *alg-4* in wild-type and *mut-16* mutant L4 animals cultured at 20°C and 25°C. Error bars indicate standard deviation. **(E)** Shown is the percentage of 22-nt and 26-nt reads mapping to *alg-3* and *alg-4* with A, T, C, or G represented in the first position of the read in wild-type L4 animals cultured at 20°C. **(F)** Shown are size profiles of all reads mapping to the *alg-3* genomic locus in wild-type and *mut-16* mutant L4 animals cultured at 20°C and 25°C. Error bars indicate standard deviation. **(G)** Shown are size profiles of all reads mapping to the *alg-4* genomic locus in wild-type and *mut-16* mutant L4 animals cultured at 20°C and 25°C. Error bars indicate standard deviation. **(H)** mRNA transcripts mapping to *alg-3* are counted and normalized to the expression level of *rpl-32*, in reads per million (RPMs), for L4 and adult wild-type and *mut-16* mutant animals cultured at 20°C and 25°C. Error bars indicate standard deviation. **(I)** mRNA transcripts mapping to *alg-4* are counted and normalized to the expression level of *rpl-32*, in reads per million (RPMs), for L4 and adult wild-type and *mut-16* mutant animals cultured at 20°C and 25°C. Error bars indicate standard deviation.

The ALG-3/4 pathway functions to maintain proper transcript levels for spermatogenesis-enriched genes to ensure thermotolerant male fertility^15^. As we saw developmental mis-coordination of spermatogenesis-enriched genes in heat stressed *mut-16* mutants, we wanted to determine whether mis-regulation of *alg-3* and *alg-4* expression across developmental stages was also occurring. To this end, we examined normalized expression profiles of the two Argonautes, as described earlier. Indeed, we found *alg-3* and *alg-4* are down-regulated during the L4 stage and then aberrantly up-regulated during adulthood in heat stressed *mut-16* mutants (Figure 6H and I, and Supplemental Figure S4D). Intriguingly, increased targeting by *mutator* complex-independent 22G-RNAs correlated with the mis-expression of *alg-3* and *alg-4* in heat stressed *mut-16* mutant adults (Figure 6H and I, and Supplementary Figure S4D and E). Here, our data suggests a model in which the *mutator* complex contributes to developmental stage-specific regulation of *alg-3* and *alg-4* expression by fine-tuning homeostatic levels of the different classes of 22G-RNAs throughout the organism’s lifetime. Proper balancing of small RNA pools across developmental time is fundamental for switching on and off *alg-3* and *alg-4*, and thus ALG-3/4 pathway function, during the correct life stage as well as enabling the animal to respond to environmental heat stress to provide thermotolerant male fertility.

Taken together, our data reveals the paternal effect of *mut-16* triggers spermiogenesis defects that underly the onset of temperature-sensitive sterility. The spermiogenesis defects manifest due to developmental mis-regulation of sperm genes, which are largely targets of the *mutator* complex-independent ALG-3/4 pathway. Previously, it was shown that disruption of the ALG-3/4 pathway in *alg-3(tm1155); alg-4(ok1041)* mutants leads to similar spermiogenesis defects and infertility at elevated temperature^15^. Here, we show proper L4 stage-specific expression of the Argonautes, ALG-3 and ALG-4, is disrupted upon heat stress in *mut-16* mutants. Our findings indicate that *mutator* complex-dependent 22G-RNA amplification is essential for maintaining homeostatic regulation of *alg-3* and *alg-4* via a small RNA-mediated genetic switch throughout development and particularly during heat stress. Thus, the *mutator* complex plays a critical role in an RNAi to RNAi-mediated cascade responsible for switching on and off ALG-3/4 pathway function and coordinating the spermatogenesis developmental program (Figure 7).

**Figure 7.**
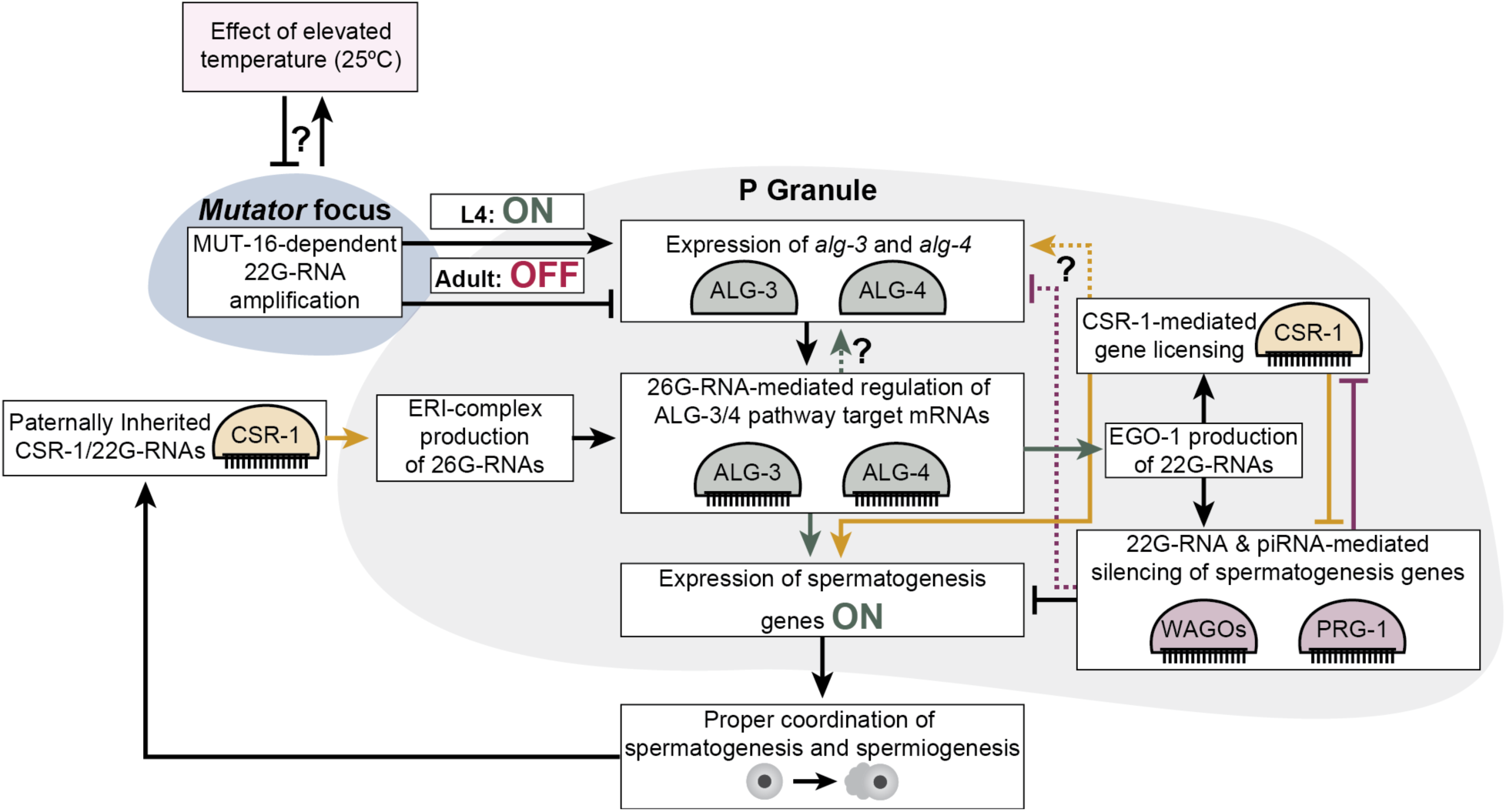
MUT-16 is required to restrict *alg-3* and *alg-4* expression and spermatogenesis to the L4 developmental stage. Model of the small RNA-mediated regulatory mechanism in which MUT-16-dependent 22G-RNAs are required to counterbalance the effects of heat stress and control expression of *alg-3* and *alg-4* to maintain developmental stage-specific ALG-3/4 pathway (green) function that, with the CSR-1 pathway (yellow) and piRNA pathway (purple), regulates expression of spermatogenesis genes to maintain sperm-based fertility. Dashed lines with question marks indicate hypothetical regulation of *alg-3* and *alg-4* by different RNAi pathways.

## DISCUSSION

Robust coordination of cellular and physiological processes throughout development is crucial for the viability of all organisms. Evolutionarily conserved RNAi pathways serve as one gene regulatory mechanism essential for proper development and maintenance of fertility in all animals. When the gene regulatory networks controlling physiological processes are disrupted, severe consequences can arise, such as loss of reproductive potential. One environmental stressor that negatively impacts gametogenesis across animal species is elevated temperature^39–41^. In *C. elegans*, many RNAi pathway mutants exhibit a transgenerational loss of fertility when exposed to elevated temperature^3, 12, 15, 24–27^. However, the maternal and paternal effects underlying temperature-sensitive sterility, and how RNAi pathways normally protect against the effects of external stressors to maintain fertility, are not fully understood.

Previously, we determined that loss of MUT-16, a factor essential for amplification of many 22G-RNAs that maintain robust heritable RNAi-mediated silencing, has both paternal and maternal effects that result in temperature-sensitive sterility after one and three generations, respectively, at elevated temperature (25°C)^24^. We characterized the maternal effect of *mut-16*, in adult animals, and found that the onset of heat stress-induced sterility correlates with a loss of germ cell identity due to aberrant expression of soma-enriched and spermatogenesis-enriched genes associated with increased germ cell chromatin accessibility at these gene loci^24^. Presently, we sought to characterize the paternal effect of *mut-16* on temperature-sensitive sterility. Through bioinformatic comparison of mRNA-seq and small RNA-seq libraries, we found that during heat stress, the paternal effect of *mut-16* triggers mis-coordination of the spermatogenesis developmental program driven by small RNA-dependent temporal mis-regulation of the Argonuates, ALG-3 and ALG-4. The developmental mis-regulation of ALG-3/4 pathway function ultimately results in spermiogenesis defects that trigger the immediate temperature-sensitive sperm-based sterility of *mut-16* mutants. These findings reveal a developmental genetic cascade, controlled by RNAi pathways and sensitive to changes in environmental temperature, is responsible for precise temporal regulation of ALG-3/4 pathway function and thus, thermotolerant male fertility.

Our findings implicate the *Mutator* focus and *mutator* complex-dependent 22G-RNA amplification in coordinating the spermatogenesis developmental program. Typically, the spermatogenesis and oogenesis programs occur sequentially during discrete developmental timeframes within the gonad of *C. elegans*. During the L4 stage, a subset of the totipotent germ cells in the gonadal arms undergo spermatogenesis to develop into spermatids, and then the remaining germ cells subsequently develop into oocytes during the adult stage. Thus, proper control of the spermatogenesis to oogenesis program switch is critical for maintaining fertility. For sperm-based fertility, transcription of spermatogenesis genes is necessary during the narrow L4 developmental timeframe but must be suppressed during the adult developmental stage for proper oocyte-based fertility. Previous works have determined that appropriate regulation of spermatogenesis genes requires the ALG-3/4, CSR-1, and piRNA pathways, which localize to P granules – perinuclear germ granules that sit adjacent to *Mutator* foci^12, 13, 15, 16, 19–23, 27, 37, 42–47^. P granules are hallmarks of germ cell identity and persist in oocytes but are disassembled in spermatids during spermiogenesis, thus their presence within the adult germline is indicative of successful completion of the spermatogenesis to oogenesis program switch^48–50^.

Here we show that the absence of MUT-16-dependent small RNAs results in reduced spermatogenesis-enriched gene expression during the L4 stage and aberrant up-regulation of sperm genes in adults. This phenomenon is observed in *mut-16* mutants cultured at permissive temperature (20°C) but is more pronounced at elevated temperature (25°C) when the animals exhibit temperature-sensitive sterility. We believe the paternal effect, and not the maternal effect, of *mut-16* underlies the delayed onset of spermatogenesis-enriched gene expression observed in adults. In addition, adult heat stressed *mut-16* mutant germlines have P granule abnormalities^24^. The aberrant expression of spermatogenesis-enriched genes, coupled with the observed P granule abnormalities, in adult germlines suggests heat stressed *mut-16* mutants exhibit a defect in the switch between the spermatogenesis and oogenesis programs. This hypothesis is also supported by the increased number of spermatids present in the *mut-16* mutant males at elevated temperature. A similar pattern of reduced spermatogenesis gene expression during the L4 developmental stage followed by sperm gene up-regulation during adulthood has been observed in *prg-1* mutants, which have a defective piRNA pathway^27, 51^. Sterility in piRNA pathway mutants also correlates with abnormalities in *Mutator* foci and P granule formation^52^. Furthermore, *csr-1* mutants, which have an aggregated P granule phenotype, and *C. elegans* lacking P granules also exhibit increased expression of spermatogenesis genes in adults resulting in masculinization of the adult hermaphroditic germline^23^. Taken together, these findings suggest that *Mutator* foci and P granules coordinate the piRNA, CSR-1 and ALG-3/4 pathways to properly initiate and terminate the spermatogenesis developmental program.

Coordination of the initiation and termination of the spermatogenesis and oogenesis developmental programs is critical for maintaining the reproductive potential of *C. elegans*. Our bioinformatic analyses revealed that loss of MUT-16-dependent 22G-RNA amplification triggers the developmental mis-regulation of the spermatogenesis program, but largely does not impact the oogenesis program. In addition, we found that expression of sperm genes is more susceptible to perturbation by heat stress than that of oogenesis genes. Our data indicates that spermatogenesis-enriched and oogenesis-enriched genes are controlled by distinct layers of regulatory mechanisms. Furthermore, our data points to the spermatogenesis and oogenesis developmental programs being independently coordinated such that initiation of one program does not rely on the termination of the other. These findings have broad implications for our understanding of the processivity and regulation of developmental programs in totipotent germ cells.

Previously, a model was proposed in which P granules and the Argonaute, CSR-1, work together to coordinate the spermatogenesis to oogenesis switch through small RNA-mediated suppression of spermatogenesis-enriched gene transcript levels in germ cells at the beginning of oogenesis^23^. This model builds on foundational work that discovered ALG-3 and ALG-4 bind 26G-RNAs complementary to sperm-enriched genes, and are expressed during spermatogenesis (L4 stage) to fine-tune the levels of sperm gene transcripts to maintain thermotolerant male fertility^15^. Many ALG-3/4 pathway and CSR-1 pathway targets overlap, so it was proposed that regulation of targets by the ALG-3/4 pathway during the L4 stage triggers MUT-16-independent 22G-RNA production from sperm gene mRNAs for downstream loading into CSR-1 and WAGO-1^16^. These CSR-1-loaded 22G-RNAs are thought to then act in a transgenerational feedforward loop providing sperm-mediated transmission of the epigenetic memory of paternal gene expression necessary for the next generation’s fertility^16^. These models relied on L4 stage-specific expression of ALG-3 and ALG-4 driving ALG-3/4 pathway function to appropriately regulate spermatogenesis-enriched genes; however, how temporal regulation of these Argonautes is achieved remained elusive.

Based on our findings, we propose that *mutator* complex-dependent 22G-RNAs play a key role in coordinating the switch from the spermatogenesis to oogenesis developmental program by controlling the expression of *alg-3* and *alg-4* (Figure 7). Our model proposes that *mutator* complex-dependent 22G-RNA amplification is required to promote the expression of *alg-3* and *alg-4* during the L4 stage, possibly in concert with autoregulatory feedback from ALG-3/4-bound 26G-RNAs. Then, during the adult stage, a change in the pool of *mutator* complex-dependent 22G-RNAs drives the genetic cascade that switches off *alg-3* and *alg-4* expression, ultimately terminating the spermatogenesis developmental program. Furthermore, our model proposes that *mutator* complex-dependent small RNA amplification is essential for mitigating the effects of heat stress and maintains robust fertility through adjusting sperm-enriched gene transcript levels in a small RNA-dependent manner to compensate for heat stress-induced molecular changes.

Our model postulates that MUT-16, acts upstream of ALG-3/4 pathway function in maintaining appropriate spermatogenesis gene expression by modulating *alg-3* and *alg-4* expression through fine-tuning of small RNAs targeting their mRNAs. If this is the case, we would expect *mut-16* mutants and *alg-3; alg-4* mutants to phenocopy each other. It was previously shown that *alg-3; alg-4* mutants exhibit ∼50% reduced fertility compared to wild-type animals at 20°C^15, 16^. During the L4 stage, we see reduced expression of *alg-3* and *alg-4* in *mut-16* mutants grown at permissive temperature (20°C) compared to wild-type L4 animals. In addition, we have shown here and previously that *mut-16* mutants have ∼50% fewer progeny compared to wild-type animals when cultured at 20°C. We think that the similar reduction in brood size at 20°C is indicative of a common defect in sperm-based fertility in *mut-16* and *alg-3; alg-4* mutants. Spermatids in *alg-3; alg-4* mutants exhibit spermiogenesis defects, specifically arresting as spiky intermediates upon activation by pronase E, and at elevated temperature (25°C) *alg-3; alg-4* mutants are sterile^15, 16^. Here, we show that heat stress exacerbates the loss of *alg-3* and *alg-4* expression in *mut-16* mutants, which correlates with increased incidence of spermatids that arrest as spiky intermediates and the onset of temperature-sensitive sterility. Taken together, these results indicate *mut-16* and *alg-3; alg-4* mutants have comparable thermosensitive sperm-based infertility phenotypes.

The ALG-3 and ALG-4 Argonautes bind 26G-RNAs, which are not synthesized by the *mutator* complex, so how does loss of *mut-16* trigger reduced 26G-RNA levels targeting spermatogenesis genes and the ALG-3/4 pathway targets? Previously, we identified a small RNA-mediated feedback loop in which MUT-16-dependent 22G-RNAs regulate the expression of the ERGO-1-class 26G-RNA biogenesis factor, ERI-6/7^38^. In this feedback mechanism, loss of MUT-16-dependent 22G-RNAs triggers reduced expression of trans-spliced *eri-6/7* mRNAs, thereby indirectly regulating the upstream production of ERGO-1-class 26G-RNAs^38^. We hypothesize the loss of 26G-RNAs targeting ALG-3/4 pathway targets, as well as *alg-3* and *alg-4*, observed in the *mut-16* mutant at elevated temperature (25°C) is the result of another small RNA-mediated feedback loop. In fact, *alg-3* and *alg-4* were previously categorized as ALG-3/4 pathway positively regulated targets; however, this was based on mRNA-seq libraries generated from *alg-3; alg-4* mutants^16^. Whether the reduction of small RNAs mapping to the genomic loci of *alg-3* and *alg-4* is the result of autoregulation by the ALG-3/4 pathway or direct regulation by one, or multiple, other RNAi pathway branches remains to be explored. Our mRNA-seq and small RNA-seq data revealed the MUT-16-dependent and MUT-16-independent small RNAs that act to regulate the expression of *alg-3* and *alg-4* are thermosensitive. This putative small RNA-mediated feedback mechanism warrants additional exploration to determine how it modulates *alg-3* and *alg-4* expression in response to changes in environmental temperature and fluctuations in small RNA populations, and to further elucidate the complexities of the RNAi pathways that regulate developmental cellular programs. Moving forward, identifying the molecular mechanism(s) underlying the thermosensitivity of small RNA populations will be critical for deepening our understanding of gene regulation as, here, we show RNAi pathways are essential for an organism’s ability to maintain proper gene expression across developmental time in the face of environmental perturbations.

Importantly, this work has identified a novel gene regulatory network architecture in which an RNAi to RNAi-mediated cascade not only controls the function of an RNAi pathway branch, but also triggers the initiation and termination of a developmental program. RNAi pathway-mediated gene regulation is critical for the viability of all animals, including humans, implying there is exciting potential for more RNAi-mediated feedback network architectures that coordinate cellular programs in eukaryotes to be discovered in the future.

## MATERIALS AND METHODS

C. elegans strains Unless otherwise stated, worms were grown at 20°C according to standard conditions^53^. All strains are in the N2 background. Strains used include:

N2 – wild-type
NL1810 – *mut-16(pk710) I. BS553 – fog-2(oz40) V*.
DUP75 – *pgl-1(sam33[pgl-1::GFP::3xFLAG]) IV*.
USC1252 – *mut-16(pk710) I; pgl-1(sam33[pgl-1::GFP::3xFLAG]) IV*.

### Male induction

L4s were plated on NGM plates and incubated at 30°C for 4, 5, and 6 hours then placed at 20°C to recover. In the broods, males were isolated and mated with young adult hermaphrodites to maintain increased incidences of males within the populations.

### Mating brood size assay

Synchronized L1s of *fog-2(oz40)*, *pgl-1::GFP::3xFLAG*, *mut-16(pk710)*, and *mut-16(pk710); pgl-1::GFP::3xFLAG* male-containing populations were plated on NGM plates and cultured at 20°C or 25°C for a single generation. Mating plates were set up using individual L4 hermaphrodites of *fog-2(oz40)* or *mut-16(pk710)* grown at 20°C or 25°C and individual males of *pgl-1::GFP::3xFLAG* or *mut-16(pk710); pgl-1::GFP::3xFLAG* grown at 20°C or 25°C (ten plates per cross per temperature). Expression of PGL-1::GFP::3xFLAG was used to distinguish cross-progeny from the hermaphrodite’s self-progeny. The number of GFP-expressing eggs laid were counted for each hermaphrodite.

### Fluorescence microscopy

Synchronized L1s of wild-type and *mut-16(pk710)* worms were plated on NGM plates and cultured at 20°C and 25°C. Males were isolated as L4s and allowed to develop to adulthood for 24 hours in the absence of hermaphrodites (to prevent mating). Whole animals were fixed in pre-chilled (−20°C) methanol for 5 minutes then washed twice in 1xPBST before being incubated in 0.17 μg/mL DAPI solution in PBST for 15 minutes, followed by three washes in 1xPBST. Imaging was performed on an AxioImager.M2 (Zeiss) using a Plan-Apochromat 63x/1.4-oil immersion objective. For sperm counting, z-stacks were processed using FIJI^54–56^ and a projection of the stack with sperm in different planes were pseudocolored using the hyperstacks temporal-color code function. The multipoint tool in FIJI was then used to count the post-meiotic sperm in each male. The stack projection was pseudo-colored using Adobe Photoshop. Post-meiotic sperm were counted for 5 biological replicates per genotype per condition.

### Spicule morphology assay

Synchronized L1s of wild-type and *mut-16(pk710)* worms were plated on NGM plates and cultured at 20°C and 25°C. Males were collected as adults (∼68hrs at 20°C and ∼48hrs at 25°C) and were live imaged. Imaging was performed on an AxioImager.M2 (Zeiss) using a Plan-Apochromat 63x/1.4-oil immersion objective. One spicule per tail was scored for spicule morphology (non-crumpled or crumpled) and spicule length. Spicule length was measured in FIJI using the line and measure tools^54–56^. Between 40 and 51 spicules were assayed per genotype per temperature condition.

### RNA Extraction

Synchronized L1s of wild-type and *mut-16(pk710)* worms were plated on enriched peptone plates and cultured at 20°C and 25°C. 8,000 animals per sample were harvested as L4s (∼56hrs at 20°C and ∼39hrs at 25°C) for RNA extraction. Worms were washed off plates using water and then settled on ice to form a pellet. Water was aspirated off and worm pellets were resuspended in 1mL TRIzol reagent (Life Technologies) and freeze-thawed on dry ice followed by vortexing. Worm carcasses were pelleted using centrifugation and the supernatant containing RNA was collected. 0.2 volume chloroform was added to supernatant, vortexed, centrifuged, and then the aqueous phase was transferred to a new tube. Samples were precipitated using isopropanol and rehydrated in 50μL nuclease-free H_2_O.

### mRNA-seq library preparation

Nuclease-free H_2_O was added to 7.5μg of each total RNA sample, extracted from whole animals to a final volume of 100μL. Samples were incubated at 65°C for 2 minutes then incubated on ice. The Dynabeads mRNA Purification Kit (ThermoFisher 61006) was used according to the manufacturer’s protocol. 20μL of Dynabeads was used for each sample. 100ng of each mRNA sample was used to prepare libraries with the NEBNext Ultra II Directional RNA Library Prep Kit for Illumina (NEB E7760S) according to the manual, using NEBNext multiplex oligos for Illumina (NEB E7335S). Library quality was assessed (Agilent BioAnalyzer Chip) and concentration was determined using the Qubit 1X dsDNA HS Assay kit (ThermoFisher Q33231). Libraries were sequenced on the Illumina NextSeq500 (SE 75-bp reads) platform. Two biological replicates were generated for wild-type (N2) and *mut-16(pk710)* mutants cultured at 20°C and 25°C.

### Small RNA library preparation

Small RNAs (18- to 30- nt) were size selected on denaturing 15% polyacrylamide gels (Criterion 3450091) from total RNA samples. Libraries were prepared as previously described^57^. Library quality was assessed (Agilent BioAnalyzer Chip) and concentration was determined using the Qubit 1X dsDNA HS Assay kit. Libraries were sequenced on the Illumina NextSeq500 (SE 75-bp reads) platform. Two biological replicates were generated for wild-type (N2) and *mut-16(pk710)* mutants cultured at 20°C and 25°C.

### Bioinformatic analysis

For small RNA libraries and mRNA libraries, sequences were parsed from adapters using Cutadapt^58^ and mapped to the *C. elegans* genome, WS258, using HISAT2^59^ and the transcriptome using Salmon^60^. Data analysis was done using R, Excel, deepTools2^61^, and custom Python scripts. Reads per million were plotted along the WS258 genome using Integrative Genomics Viewer 2.3.68^62^. Tissue enrichment analysis was performed using reference gene lists from WormExp^63^. CSR-1 target genes, ALG-3/4 target genes, ERGO-1 target genes, *mutator* target genes, piRNA target genes, spermatogenesis-enriched genes, oogenesis-enriched genes, soma-enriched genes, muscle-enriched genes, and neuron-enriched genes were previously described^12, 14, 16, 42, 64–69^. Sequencing data is summarized in Supplementary Table 1.

### cDNA preparation and qPCR reactions

RNA samples were DNAse treated using DNAse I, Amplification Grade (Invitrogen 18068015), and reverse transcribed with SuperScript IV Reverse Transcriptase (Invitrogen 18090050), following manufacturers’ protocols. All Real time PCR reactions were performed using the PowerTrack SYBR Green Master Mix (Applied Biosystems A46109) and run on the QuantStudio 3 Real-Time PCR System (Applied Biosystems A28567). Primers used are listed in Supplementary Table 2.

### *In vitro* sperm activation assay

L4 males of wild-type and *mut-16(pk710)* worms were isolated on NGM plates without hermaphrodites and allowed to develop into adults at 20°C and 25°C for 24 hours. 10-15 males were dissected in 15 μL sperm buffer (50mM HEPES, 50 mM NaCl, 25mM KCl, 5 mM CaCl_2_, 1 mM MgSO_4_, 0.1% BSA) with or without 400 ng/mL Pronase E (Millipore Sigma P8811) as described previously^70^. Tails were nicked off to release the sperm, and the slides were incubated in a hybridization chamber for 15 minutes before a coverslip was mounted. Sperm were imaged on an AxioImager.M2 (Zeiss) using a Plan-Apochromat 63x/1.4-oil immersion objective. Pseudopod formation was scored for 200 sperm per genotype per temperature condition. Pseudopod formation data is summarized in Supplementary Table 3.

## DATA AVAILABILITY

High-throughput sequencing data for mRNA-seq and small RNA-seq experiments generated during this study are available through Gene Expression Omnibus (GEO #: GSE226893). De-multiplexed raw sequencing data, in fastq format, for N2 and *mut-16(pk710)* adult mRNA-seq and small RNA-seq libraries used from^24^ were obtained from NCBI’s Gene Expression Omnibus (GEO #: GSE134573).

## SUPPLEMENTARY DATA

Supplementary Data are available online.

## ACKNOWLEDGEMENT

We thank the members of the Rogers lab, Mark Pellegrino, Scott H. Saunders, and Carolyn M. Phillips for helpful discussions and feedback on the manuscript. Some strains were provided by the CGC, which is funded by NIH Office of Research Infrastructure Programs (P40 OD010440). Next generation sequencing was performed by the USC Molecular Genomics Core, which is supported by award number P30 CA014089 from the National Cancer Institute.

## AUTHOR CONTRIBUTIONS

Conceptualization, A.K.R.; Investigation, A.K.R. and C.M.; Formal Analysis, A.K.R.; Writing, A.K.R.; Visualization, A.K.R.; Funding Acquisition, A.K.R.; Supervision, A.K.R.

## REFERENCES

1. Riddle, D.L., Blumenthal, T., Meyer, B.J., and Priess, J.R. (1997). C. elegans II (Cold Spring Harbor Laboratory Press).

2. Klass, M., Wolf, N., and Hirsh, D. (1976). Development of the male reproductive system and sexual transformation in the nematode Caenorhabditis elegans. Dev Biol 52, 1–18. 10.1016/0012-1606(76)90002-6.

3. Buckley, B.A., Burkhart, K.B., Gu, S.G., Spracklin, G., Kershner, A., Fritz, H., Kimble, J., Fire, A., and Kennedy, S. (2012). A nuclear Argonaute promotes multigenerational epigenetic inheritance and germline immortality. Nature 489, 447–451. 10.1038/nature11352.

4. Burkhart, K.B., Guang, S., Buckley, B.A., Wong, L., Bochner, A.F., and Kennedy, S. (2011). A pre-mRNA-associating factor links endogenous siRNAs to chromatin regulation. PLoS Genet 7, e1002249. 10.1371/journal.pgen.1002249.

5. Claycomb, J.M. (2014). Ancient endo-siRNA pathways reveal new tricks. Curr Biol 24, R703–715. 10.1016/j.cub.2014.06.009.

6. Gu, S.G., Pak, J., Guang, S., Maniar, J.M., Kennedy, S., and Fire, A. (2012). Amplification of siRNA in Caenorhabditis elegans generates a transgenerational sequence-targeted histone H3 lysine 9 methylation footprint. Nat Genet 44, 157–164. 10.1038/ng.1039.

7. Guang, S., Bochner, A.F., Pavelec, D.M., Burkhart, K.B., Harding, S., Lachowiec, J., and Kennedy, S. (2008). An Argonaute transports siRNAs from the cytoplasm to the nucleus. Science 321, 537–541. 10.1126/science.1157647.

8. Guang, S., Bochner, A.F., Burkhart, K.B., Burton, N., Pavelec, D.M., and Kennedy, S. (2010). Small regulatory RNAs inhibit RNA polymerase II during the elongation phase of transcription. Nature 465, 1097–1101. 10.1038/nature09095.

9. Hutvagner, G., and Simard, M.J. (2008). Argonaute proteins: key players in RNA silencing. Nat Rev Mol Cell Biol 9, 22–32. 10.1038/nrm2321.

10. Bernstein, E., Caudy, A.A., Hammond, S.M., and Hannon, G.J. (2001). Role for a bidentate ribonuclease in the initiation step of RNA interference. Nature 409, 363–366. 10.1038/35053110.

11. Ketting, R.F., Fischer, S.E., Bernstein, E., Sijen, T., Hannon, G.J., and Plasterk, R.H. (2001). Dicer functions in RNA interference and in synthesis of small RNA involved in developmental timing in C. elegans. Genes Dev 15, 2654–2659. 10.1101/gad.927801.

12. Gu, W., Shirayama, M., Conte, D., Vasale, J., Batista, P.J., Claycomb, J.M., Moresco, J.J., Youngman, E.M., Keys, J., Stoltz, M.J., et al. (2009). Distinct argonaute-mediated 22G-RNA pathways direct genome surveillance in the C. elegans germline. Mol Cell 36, 231–244. 10.1016/j.molcel.2009.09.020.

13. Claycomb, J.M., Batista, P.J., Pang, K.M., Gu, W., Vasale, J.J., van Wolfswinkel, J.C., Chaves, D.A., Shirayama, M., Mitani, S., Ketting, R.F., et al. (2009). The Argonaute CSR-1 and its 22G-RNA cofactors are required for holocentric chromosome segregation. Cell 139, 123–134. 10.1016/j.cell.2009.09.014.

14. Zhang, C., Montgomery, T.A., Gabel, H.W., Fischer, S.E., Phillips, C.M., Fahlgren, N., Sullivan, C.M., Carrington, J.C., and Ruvkun, G. (2011). mut-16 and other mutator class genes modulate 22G and 26G siRNA pathways in Caenorhabditis elegans. Proc Natl Acad Sci U S A 108, 1201–1208. 10.1073/pnas.1018695108.

15. Conine, C.C., Batista, P.J., Gu, W., Claycomb, J.M., Chaves, D.A., Shirayama, M., and Mello, C.C. (2010). Argonautes ALG-3 and ALG-4 are required for spermatogenesis-specific 26G-RNAs and thermotolerant sperm in Caenorhabditis elegans. Proc Natl Acad Sci U S A 107, 3588–3593. 10.1073/pnas.0911685107.

16. Conine, C.C., Moresco, J.J., Gu, W., Shirayama, M., Conte, D., Jr., Yates, J.R., 3rd, and Mello, C.C. (2013). Argonautes promote male fertility and provide a paternal memory of germline gene expression in C. elegans. Cell 155, 1532–1544. 10.1016/j.cell.2013.11.032.

17. Han, T., Manoharan, A.P., Harkins, T.T., Bouffard, P., Fitzpatrick, C., Chu, D.S., Thierry-Mieg, D., Thierry-Mieg, J., and Kim, J.K. (2009). 26G endo-siRNAs regulate spermatogenic and zygotic gene expression in Caenorhabditis elegans. Proc Natl Acad Sci U S A 106, 18674–18679. 10.1073/pnas.0906378106.

18. Pavelec, D.M., Lachowiec, J., Duchaine, T.F., Smith, H.E., and Kennedy, S. (2009). Requirement for the ERI/DICER complex in endogenous RNA interference and sperm development in Caenorhabditis elegans. Genetics 183, 1283–1295. 10.1534/genetics.109.108134.

19. Seth, M., Shirayama, M., Gu, W., Ishidate, T., Conte, D., and Mello, C.C. (2013). The C. elegans CSR-1 argonaute pathway counteracts epigenetic silencing to promote germline gene expression. Dev Cell 27, 656–663. 10.1016/j.devcel.2013.11.014.

20. Wedeles, C.J., Wu, M.Z., and Claycomb, J.M. (2013). Protection of germline gene expression by the C. elegans Argonaute CSR-1. Dev Cell 27, 664–671. 10.1016/j.devcel.2013.11.016.

21. Nguyen, D.A.H., and Phillips, C.M. (2021). Arginine methylation promotes siRNA-binding specificity for a spermatogenesis-specific isoform of the Argonaute protein CSR-1. Nat Commun 12, 4212. 10.1038/s41467-021-24526-6.

22. Charlesworth, A.G., Seroussi, U., Lehrbach, N.J., Renaud, M.S., Sundby, A.E., Molnar, R.I., Lao, R.X., Willis, A.R., Woock, J.R., Aber, M.J., et al. (2021). Two isoforms of the essential C. elegans Argonaute CSR-1 differentially regulate sperm and oocyte fertility. Nucleic Acids Res 49, 8836–8865. 10.1093/nar/gkab619.

23. Campbell, A.C., and Updike, D.L. (2015). CSR-1 and P granules suppress sperm-specific transcription in the C. elegans germline. Development 142, 1745–1755. 10.1242/dev.121434.

24. Rogers, A.K., and Phillips, C.M. (2020). RNAi pathways repress reprogramming of C. elegans germ cells during heat stress. Nucleic Acids Res 48, 4256–4273. 10.1093/nar/gkaa174.

25. Ni, J.Z., Kalinava, N., Chen, E., Huang, A., Trinh, T., and Gu, S.G. (2016). A transgenerational role of the germline nuclear RNAi pathway in repressing heat stress-induced transcriptional activation in C. elegans. Epigenetics Chromatin 9, 3. 10.1186/s13072-016-0052-x.

26. Manage, K.I., Rogers, A.K., Wallis, D.C., Uebel, C.J., Anderson, D.C., Nguyen, D.A.H., Arca, K., Brown, K.C., Cordeiro Rodrigues, R.J., de Albuquerque, B.F., et al. (2020). A tudor domain protein, SIMR-1, promotes siRNA production at piRNA-targeted mRNAs in. Elife 9. 10.7554/eLife.56731.

27. Wang, G., and Reinke, V. (2008). A C. elegans Piwi, PRG-1, regulates 21U-RNAs during spermatogenesis. Curr Biol 18, 861–867. 10.1016/j.cub.2008.05.009.

28. Schedl, T., and Kimble, J. (1988). fog-2, a germ-line-specific sex determination gene required for hermaphrodite spermatogenesis in Caenorhabditis elegans. Genetics 119, 43–61. 10.1093/genetics/119.1.43.

29. Nett, E.M., Sepulveda, N.B., and Petrella, L.N. (2019). Defects in mating behavior and tail morphology are the primary cause of sterility in. J Exp Biol 222. 10.1242/jeb.208041.

30. L’Hernault, S.W. (2006). Spermatogenesis. WormBook, 1–14. 10.1895/wormbook.1.85.1.

31. Shakes, D.C., and Ward, S. (1989). Initiation of spermiogenesis in C. elegans: a pharmacological and genetic analysis. Dev Biol 134, 189–200. 10.1016/0012-1606(89)90088-2.

32. Achanzar, W.E., and Ward, S. (1997). A nematode gene required for sperm vesicle fusion. J Cell Sci 110 (Pt 9), 1073–1081. 10.1242/jcs.110.9.1073.

33. Roberts, T.M., and Ward, S. (1982). Centripetal flow of pseudopodial surface components could propel the amoeboid movement of Caenorhabditis elegans spermatozoa. J Cell Biol 92, 132–138. 10.1083/jcb.92.1.132.

34. Ward, S., Argon, Y., and Nelson, G.A. (1981). Sperm morphogenesis in wild-type and fertilization-defective mutants of Caenorhabditis elegans. J Cell Biol 91, 26–44. 10.1083/jcb.91.1.26.

35. Washington, N.L., and Ward, S. (2006). FER-1 regulates Ca2+ -mediated membrane fusion during C. elegans spermatogenesis. J Cell Sci 119, 2552–2562. 10.1242/jcs.02980.

36. Stanfield, G.M., and Villeneuve, A.M. (2006). Regulation of sperm activation by SWM-1 is required for reproductive success of C. elegans males. Curr Biol 16, 252–263. 10.1016/j.cub.2005.12.041.

37. Phillips, C.M., Montgomery, T.A., Breen, P.C., and Ruvkun, G. (2012). MUT-16 promotes formation of perinuclear mutator foci required for RNA silencing in the C. elegans germline. Genes Dev 26, 1433–1444. 10.1101/gad.193904.112.

38. Rogers, A.K., and Phillips, C.M. (2020). A Small-RNA-Mediated Feedback Loop Maintains Proper Levels of 22G-RNAs in C. elegans. Cell Rep 33, 108279. 10.1016/j.celrep.2020.108279.

39. Harvey, S.C., and Viney, M.E. (2007). Thermal variation reveals natural variation between isolates of Caenorhabditis elegans. J Exp Zool B Mol Dev Evol 308, 409–416. 10.1002/jez.b.21161.

40. Wang, C., Cui, Y.G., Wang, X.H., Jia, Y., Sinha Hikim, A., Lue, Y.H., Tong, J.S., Qian, L.X., Sha, J.H., Zhou, Z.M., et al. (2007). transient scrotal hyperthermia and levonorgestrel enhance testosterone-induced spermatogenesis suppression in men through increased germ cell apoptosis. J Clin Endocrinol Metab 92, 3292–3304. 10.1210/jc.2007-0367.

41. Rohmer, C., David, J.R., Moreteau, B., and Joly, D. (2004). Heat induced male sterility in Drosophila melanogaster: adaptive genetic variations among geographic populations and role of the Y chromosome. J Exp Biol 207, 2735–2743. 10.1242/jeb.01087.

42. Lee, H.C., Gu, W., Shirayama, M., Youngman, E., Conte, D., and Mello, C.C. (2012). C. elegans piRNAs mediate the genome-wide surveillance of germline transcripts. Cell 150, 78–87. 10.1016/j.cell.2012.06.016.

43. Shirayama, M., Seth, M., Lee, H.C., Gu, W., Ishidate, T., Conte, D., and Mello, C.C. (2012). piRNAs initiate an epigenetic memory of nonself RNA in the C. elegans germline. Cell 150, 65–77. 10.1016/j.cell.2012.06.015.

44. Cecere, G., Hoersch, S., O’Keeffe, S., Sachidanandam, R., and Grishok, A. (2014). Global effects of the CSR-1 RNA interference pathway on the transcriptional landscape. Nature Structural & Molecular Biology 21, 358–365. doi:10.1038/nsmb.2801.

45. Updike, D., and Strome, S. (2010). P granule assembly and function in Caenorhabditis elegans germ cells. J Androl 31, 53–60. 10.2164/jandrol.109.008292.

46. Updike, D.L., Knutson, A.K., Egelhofer, T.A., Campbell, A.C., and Strome, S. (2014). Germ-granule components prevent somatic development in the C. elegans germline. Curr Biol 24, 970–975. 10.1016/j.cub.2014.03.015.

47. Uebel, C.J., Agbede, D., Wallis, D.C., and Phillips, C.M. (2020). Foci Are Regulated by Developmental Stage, RNA, and the Germline Cell Cycle in. G3 (Bethesda) 10, 3719–3728. 10.1534/g3.120.401514.

48. Amiri, A., Keiper, B.D., Kawasaki, I., Fan, Y., Kohara, Y., Rhoads, R.E., and Strome, S. (2001). An isoform of eIF4E is a component of germ granules and is required for spermatogenesis in C. elegans. Development 128, 3899–3912. 10.1242/dev.128.20.3899.

49. Kawasaki, I., Shim, Y.H., Kirchner, J., Kaminker, J., Wood, W.B., and Strome, S. (1998). PGL-1, a predicted RNA-binding component of germ granules, is essential for fertility in C. elegans. Cell 94, 635–645. 10.1016/s0092-8674(00)81605-0.

50. Kawasaki, I., Amiri, A., Fan, Y., Meyer, N., Dunkelbarger, S., Motohashi, T., Karashima, T., Bossinger, O., and Strome, S. (2004). The PGL family proteins associate with germ granules and function redundantly in Caenorhabditis elegans germline development. Genetics 167, 645–661. 10.1534/genetics.103.023093.

51. Reed, K.J., Department of Biology, C.S.U., Fort Collins, CO 80523, USA, Cell and Molecular Biology Program, C.S.U., Fort Collins, CO 80523, USA, Svendsen, J.M., Department of Biology, C.S.U., Fort Collins, CO 80523, USA, Cell and Molecular Biology Program, C.S.U., Fort Collins, CO 80523, USA, Brown, K.C., Department of Biology, C.S.U., Fort Collins, CO 80523, USA, Cell and Molecular Biology Program, C.S.U., Fort Collins, CO 80523, USA, Montgomery, B.E., et al. (2023). Widespread roles for piRNAs and WAGO-class siRNAs in shaping the germline transcriptome of Caenorhabditis elegans. Nucleic Acids Research 48, 1811–1827. 10.1093/nar/gkz1178.

52. Spichal, M., Heestand, B., Billmyre, K.K., Frenk, S., Mello, C.C., and Ahmed, S. (2021). Germ granule dysfunction is a hallmark and mirror of Piwi mutant sterility. Nat Commun 12, 1420. 10.1038/s41467-021-21635-0.

53. Brenner, S. (1974). The genetics of Caenorhabditis elegans. Genetics 77, 71–94. 10.1093/genetics/77.1.71.

54. Schindelin, J., Arganda-Carreras, I., Frise, E., Kaynig, V., Longair, M., Pietzsch, T., Preibisch, S., Rueden, C., Saalfeld, S., Schmid, B., et al. (2012). Fiji: an open-source platform for biological-image analysis. Nat Methods 9, 676–682. 10.1038/nmeth.2019.

55. Rueden, C.T., Schindelin, J., Hiner, M.C., DeZonia, B.E., Walter, A.E., Arena, E.T., and Eliceiri, K.W. (2017). ImageJ2: ImageJ for the next generation of scientific image data. BMC Bioinformatics 18, 529. 10.1186/s12859-017-1934-z.

56. Schneider, C.A., Rasband, W.S., and Eliceiri, K.W. (2012). NIH Image to ImageJ: 25 years of image analysis. Nat Methods 9, 671–675. 10.1038/nmeth.2089.

57. Montgomery, T.A., Rim, Y.S., Zhang, C., Dowen, R.H., Phillips, C.M., Fischer, S.E., and Ruvkun, G. (2012). PIWI associated siRNAs and piRNAs specifically require the Caenorhabditis elegans HEN1 ortholog henn-1. PLoS Genet 8, e1002616. 10.1371/journal.pgen.1002616.

58. Martin, M. (2011). Cutadapt removes adapter sequences from high-throughput sequencing reads. EMBnet.journal 17, 10.10.14806/ej.17.1.200.

59. Kim, D., Langmead, B., and Salzberg, S.L. (2015). HISAT: a fast spliced aligner with low memory requirements. Nat Methods 12, 357–360. 10.1038/nmeth.3317.

60. Patro, R., Duggal, G., Love, M.I., Irizarry, R.A., and Kingsford, C. (2017). Salmon provides fast and bias-aware quantification of transcript expression. Nat Methods 14, 417–419. 10.1038/nmeth.4197.

61. Ramírez, F., Ryan, D.P., Grüning, B., Bhardwaj, V., Kilpert, F., Richter, A.S., Heyne, S., Dündar, F., and Manke, T. (2016). deepTools2: a next generation web server for deep-sequencing data analysis. Nucleic Acids Res 44, W160–165. 10.1093/nar/gkw257.

62. Robinson, J.T., Thorvaldsdóttir, H., Winckler, W., Guttman, M., Lander, E.S., Getz, G., and Mesirov, J.P. (2011). Integrative genomics viewer. Nat Biotechnol 29, 24–26. 10.1038/nbt.1754.

63. Yang, W., Dierking, K., and Schulenburg, H. (2016). WormExp: a web-based application for a Caenorhabditis elegans-specific gene expression enrichment analysis. Bioinformatics 32, 943–945. 10.1093/bioinformatics/btv667.

64. Fischer, S.E., Montgomery, T.A., Zhang, C., Fahlgren, N., Breen, P.C., Hwang, A., Sullivan, C.M., Carrington, J.C., and Ruvkun, G. (2011). The ERI-6/7 helicase acts at the first stage of an siRNA amplification pathway that targets recent gene duplications. PLoS Genet 7, e1002369. 10.1371/journal.pgen.1002369.

65. Pauli, F., Liu, Y., Kim, Y.A., Chen, P.-J., and Kim, S.K. (2006). Chromosomal clustering and GATA transcriptional regulation ofintestine-expressed genes in C. elegans. 10.1242/dev.02185.

66. Watson, J.D., Wang, S., Von Stetina, S.E., Spencer, W.C., Levy, S., Dexheimer, P.J., Kurn, N., Heath, J.D., and Miller, D.M., 3rd (2008). Complementary RNA amplification methods enhance microarray identification of transcripts expressed in the C. elegans nervous system. BMC Genomics 9, 84. 10.1186/1471-2164-9-84.

67. Phillips, C.M., Montgomery, B.E., Breen, P.C., Roovers, E.F., Rim, Y.S., Ohsumi, T.K., Newman, M.A., van Wolfswinkel, J.C., Ketting, R.F., Ruvkun, G., and Montgomery, T.A. (2014). MUT-14 and SMUT-1 DEAD box RNA helicases have overlapping roles in germline RNAi and endogenous siRNA formation. Curr Biol 24, 839–844. 10.1016/j.cub.2014.02.060.

68. Tsai, H.Y., Chen, C.C., Conte, D., Jr., Moresco, J.J., Chaves, D.A., Mitani, S., Yates, J.R., 3rd, Tsai, M.D., and Mello, C.C. (2015). A ribonuclease coordinates siRNA amplification and mRNA cleavage during RNAi. Cell 160, 407–419. 10.1016/j.cell.2015.01.010.

69. Reinke, V., Gil, I.S., Ward, S., and Kazmer, K. (2004). Genome-wide germline-enriched and sex-biased expression profiles in Caenorhabditis elegans. Development 131, 311–323. 10.1242/dev.00914.

70. Hammerquist, A.M., Yen, C.A., and Curran, S.P. (2021). Analysis of *Caenorhabditis elegans* Sperm Number, Size, Activation, and Mitochondrial Content. Bio Protoc 11, e4035. 10.21769/BioProtoc.4035.

