## Supplementary Data and Tables for "RNAi-mediated regulation of *alg-3* and *alg-4* coordinates the spermatogenesis developmental program in *C. elegans*"

#### Supplementary Figures and Figure Legends

##### SUPPLEMENTARY FIGURE S1

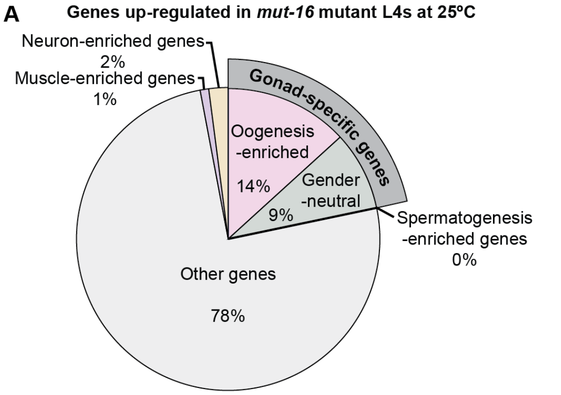


**Enrichment analysis of genes up-regulated in *mut-16* mutants at 25ºC**

| Gene lists | $\log_2$(enrichment) | Significance |
|---|---|---|
| Spermatogenesis-enriched genes | -INF | *** |
| Oogenesis-enriched genes | 2.64 | **** |
| Gender-neutral gonad genes | 1.72 | **** |
| Muscle & neuronal-specific genes | -0.33 | n.s. |

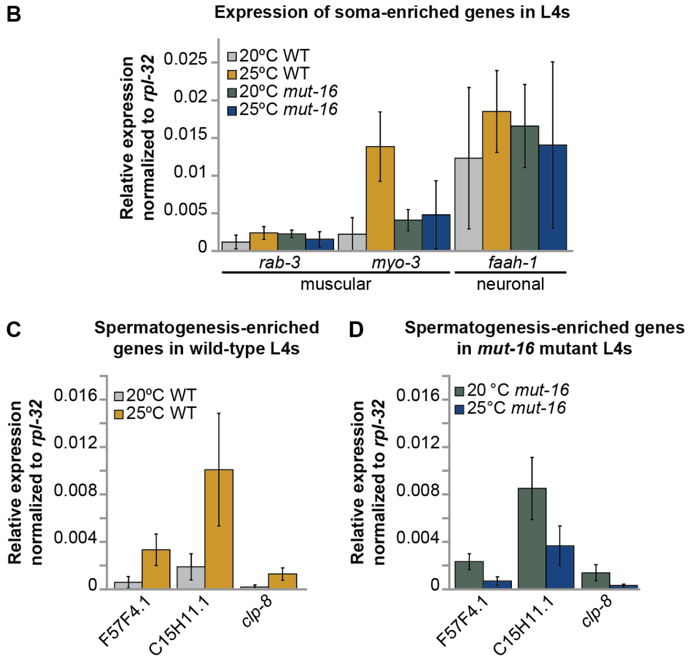


**Supplementary Figure S1. (A)** Percentages of gonad-specific and non-gonad-specific genes represented in the genes up-regulated exclusively in *mut-16* mutant L4s at 25°C, compared to wild-type L4 animals at 20°C and 25°C and *mut-16* mutant L4s at 20°C. Enrichment analysis for spermatogenesis, oogenesis, gender-neutral genes, and muscle-specific and neuronal-specific genes amongst the genes up-regulated during heat stress in *mut-16* mutants is in the table below. **(B)** qRT-PCR for genes expressed in muscle and neuronal cells in wild-type and *mut-16* mutant L4 animals grown at 20ºC and 25ºC. Expression is normalized to *rpl-32*. Error bars indicate standard deviation. n = 4 biological replicates. **(C)** qRT-PCR of spermatogenesis-enriched genes (previously observed to be up-regulated in the germline of adult *mut-16* mutants at 25ºC[24]) in wild-type L4 animals grown at 20ºC and 25ºC. Expression is normalized to *rpl-32*. Error bars indicate standard deviation. n = 4 biological replicates. **(D)** qRT-PCR of spermatogenesis-enriched genes (previously observed to be up-regulated in the germline of adult *mut-16* mutants at 25ºC[24]) in *mut-16* mutant L4 animals grown at 20ºC and 25ºC. Expression is normalized to *rpl-32*. Error bars indicate standard deviation. n = 4 biological replicates. n.s. denotes not significant and indicates a p-value > 0.05, *** indicates a p-value ≤ 0.001, and **** indicates a p-value ≤ 0.0001.

### SUPPLEMENTARY FIGURE S2

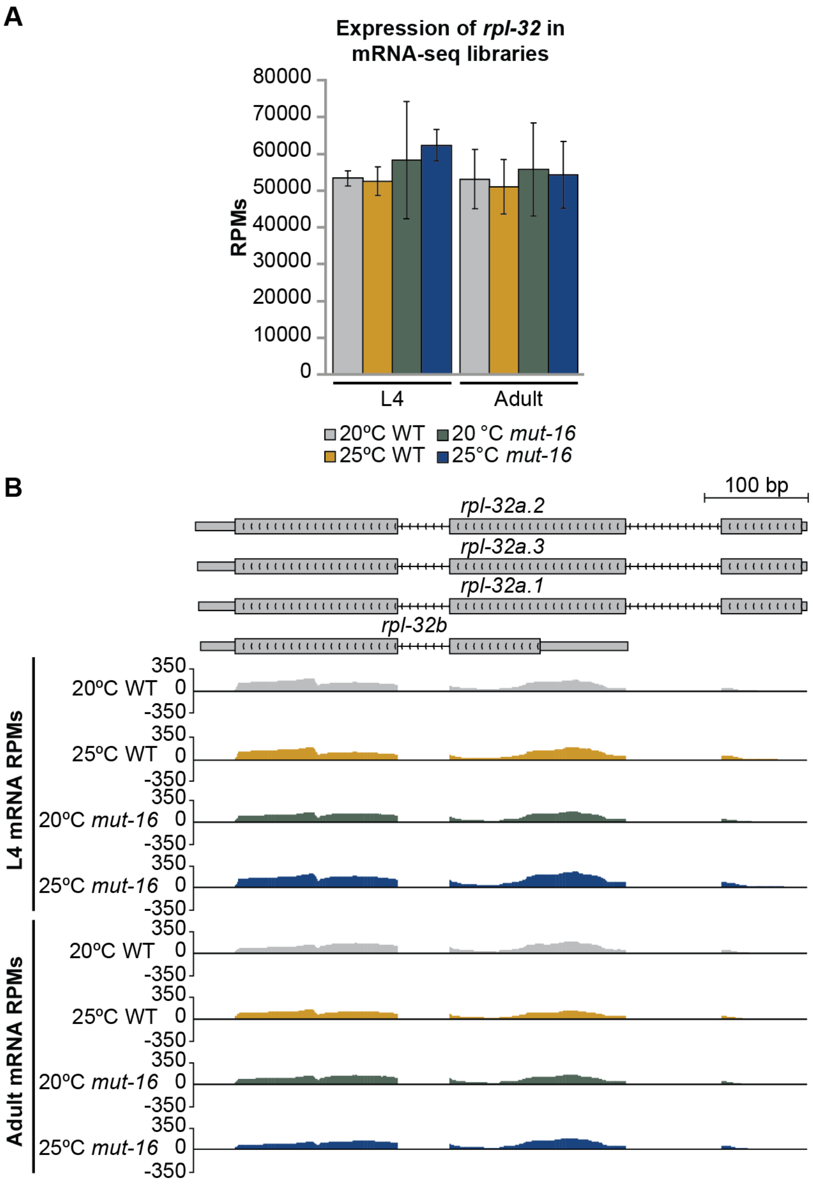


**Supplementary Figure S2. (A)** mRNA reads mapping to *rpl-32* are counted, in reads per million (RPMs), for wild-type and *mut-16* mutant L4 and adult animals cultured at 20°C and 25°C. Error bars indicate standard deviation. **(B)** Representative track of mRNA reads per million (RPMs) mapping to the *rpl-32* genomic locus in wild-type and *mut-16* mutant L4 and adult animals cultured at 20°C and 25°C.

#### SUPPLEMENTARY FIGURE S3

A

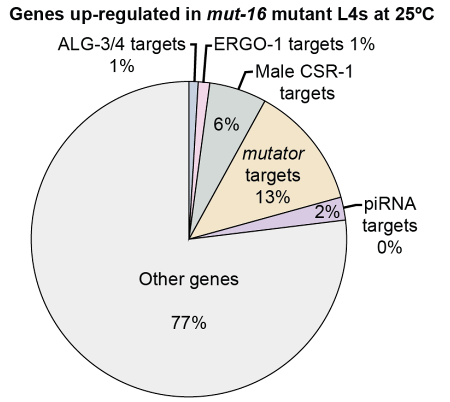


**Enrichment analysis of small RNA targets for genes up-regulated in *mut-16* mutants at 25ºC**

| Gene lists | $log_2$(enrichment) | Significance |
|---|---|---|
| ALG-3/4 targets | -1.69 | ** |
| ERGO-1 targets | 0.92 | n.s. |
| Male CSR-1 targets | -0.40 | n.s. |
| *mutator* targets | 1.57 | **** |
| piRNA targets | 2.04 | *** |

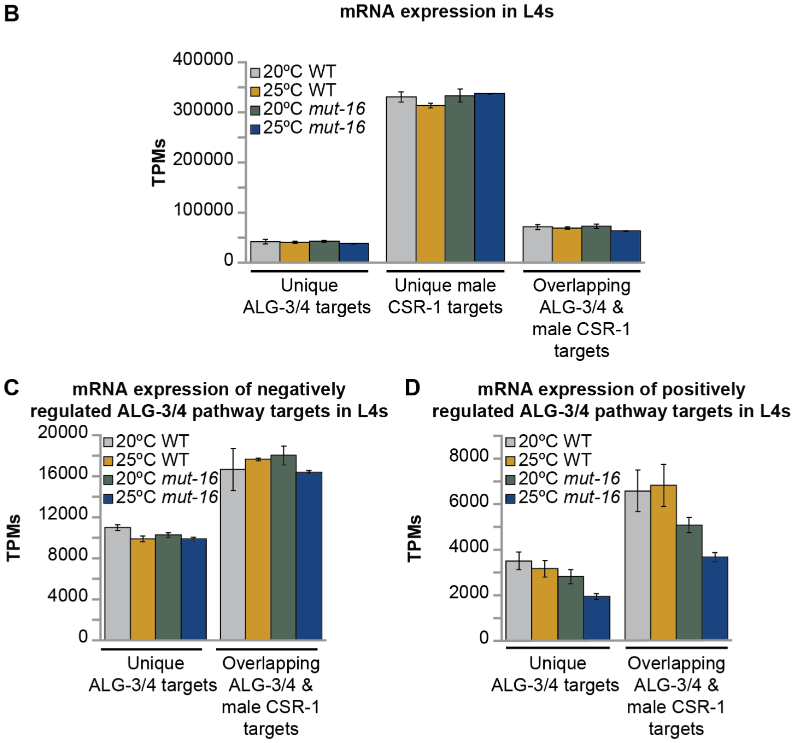


**Supplementary Figure S3. (A)** Percentages of genes targeted by distinct small RNA pathways (ALG-3/4, ERGO-1, male CSR-1, *mutator*, and piRNA pathways) represented in the genes up-regulated exclusively in *mut-16* mutant L4s at 25°C, compared to wild-type L4 animals at 20°C and 25°C and *mut-16* mutant L4s at 20°C. Enrichment analysis for ALG-3/4, ERGO-1, male CSR-1, *mutator*, and piRNA pathway targets amongst the genes up-regulated during heat stress in *mut-16* mutants is in the table below. **(B)** Shown in transcripts per million (TPMs) are mRNA transcripts mapping to genes uniquely targeted by the ALG-3/4 pathway, uniquely targeted by the male CSR-1 pathway, or genes targeted by both the ALG-3/4 and male CSR-1 pathway for wild-type and *mut-16* mutant L4 animals cultured at 20°C and 25°C. Error bars indicate standard deviation. **(C)** Shown in transcripts per million (TPMs) are mRNA transcripts mapping to negatively regulated ALG-3/4 targets that are uniquely targeted by the ALG-3/4 pathway or also targeted by the male CSR-1 pathway for wild-type and *mut-16* mutant L4 animals cultured at 20°C and 25°C. Error bars indicate standard deviation. **(D)** Shown in transcripts per million (TPMs) are mRNA transcripts mapping to positively regulated ALG-3/4 targets that are uniquely targeted by the ALG-3/4 pathway or also targeted by the male CSR-1 pathway for wild-type and *mut-16* mutant L4 animals cultured at 20°C and 25°C. Error bars indicate standard deviation.

#### SUPPLEMENTARY FIGURE S4

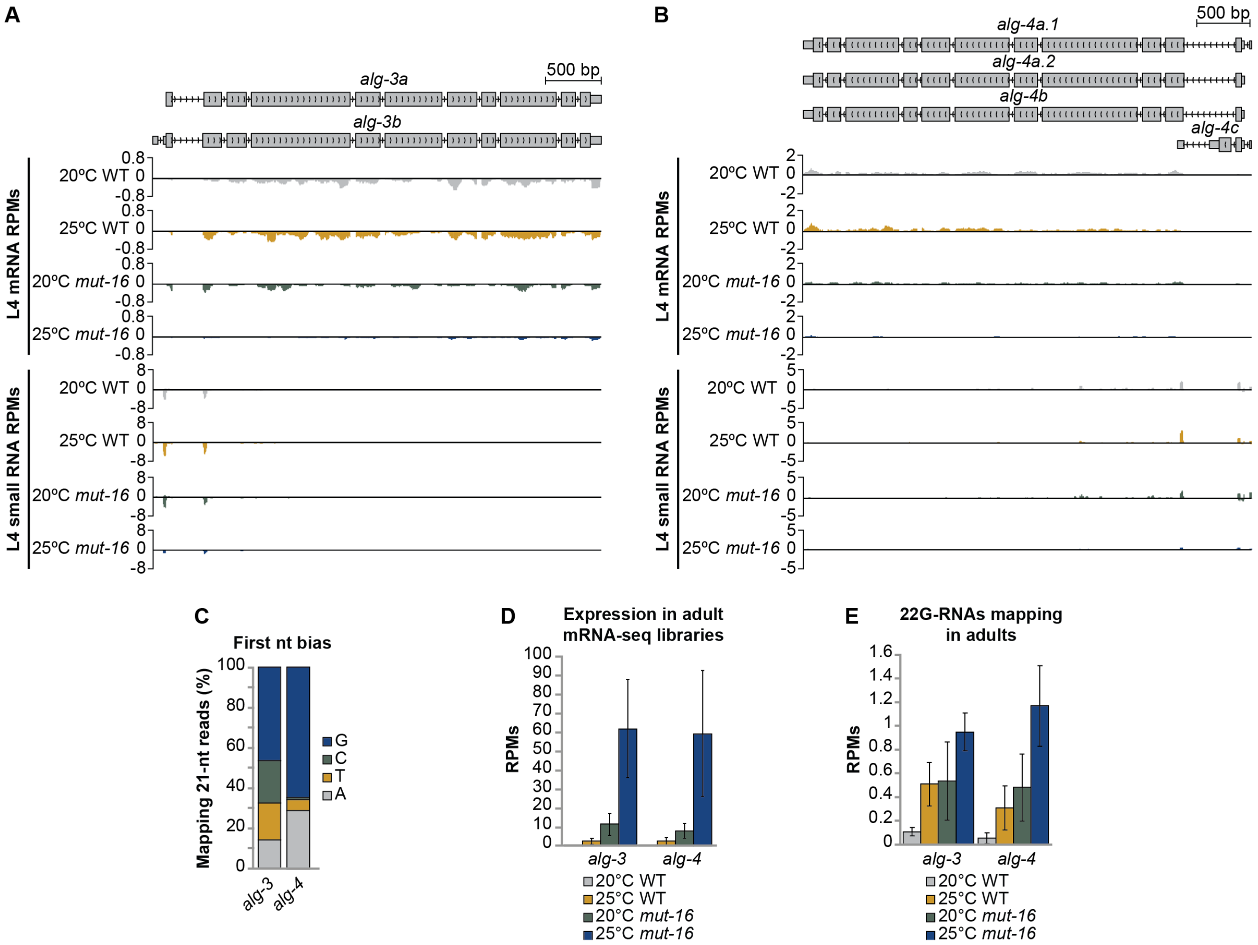


**Supplementary Figure S4. (A)** Representative tracks shown for mRNA and small RNA reads per million (RPMs) mapping to the *alg-3* genomic locus in wild-type and *mut-16* mutant L4 animals cultured at 20°C and 25°C. **(B)** Representative tracks shown for mRNA and small RNA reads per million (RPMs) mapping to the *alg-4* genomic locus in wild-type and *mut-16* mutant L4 and adult animals cultured at 20°C and 25°C. **(C)** Shown is the percentage of 21-nt reads mapping to *alg-3* and *alg-4* with A, T, C, or G represented in the first position of the read in wild-type L4 animals cultured at 20ºC. **(D)** mRNA transcripts mapping to *alg-3* and *alg-4* are counted, in reads per million (RPMs), for wild-type and *mut-16* mutant adult animals cultured at 20°C and 25°C. Error bars indicate standard deviation. **(E)** 22G-RNAs mapping to *alg-3* and *alg-4* are counted, in reads per million (RPMs), for wild-type and *mut-16* mutant adult animals cultured at 20°C and 25°C. Error bars indicate standard deviation.

**Supplementary Tables**

**Table S1. Library mapping statistics.** Related to mRNA-seq & small RNA-seq experiments in Methods.

| **Library** | **Total number of reads** | **Reads mapping to WS258 genome** | **Reads mapping to WS258 mRNA, ncRNA, and pseudogenic transcripts** |
|---|---|---|---|
| L4_N2_20C_1_mRNA | 31,939,154 | 31,205,387 | 12,178,652 |
| L4_N2_20C_2_mRNA | 30,602,733 | 29,929,582 | 10,813,181 |
| L4_*mut-16*_20C_1_mRNA | 24,731,385 | 24,088,431 | 9,914,053 |
| L4_*mut-16*_20C_2_mRNA | 30,619,535 | 29,908,181 | 12,552,697 |
| L4_N2_25C_1_mRNA | 33,666,511 | 32,906,790 | 10,997,326 |
| L4_N2_25C_2_mRNA | 35,045,891 | 34,206,941 | 12,259,241 |
| L4_*mut-16*_25C_1_mRNA | 33,728,292 | 32,980,606 | 12,276,552 |
| L4_*mut-16*_25C_2_mRNA | 33,333,928 | 32,621,587 | 13,227,929 |
| L4_N2_20C_1_small_RNA | 15,614,472 | 13,455,532 | 5,028,484 |
| L4_N2_20C_2_ small_RNA | 20,125,845 | 17,400,105 | 6,766,449 |
| L4_*mut-16*_20C_1_ small_RNA | 12,469,506 | 10,616,417 | 1,999,811 |
| L4_*mut-16*_20C_2_ small_RNA | 16,087,139 | 13,679,596 | 2,687,301 |
| L4_N2_25C_1_ small_RNA | 15,716,559 | 13,559,035 | 4,826,359 |
| L4_N2_25C_2_ small_RNA | 14,611,216 | 12,635,423 | 4,540,738 |
| L4_*mut-16*_25C_1_ small_RNA | 12,332,587 | 10,427,740 | 2,140,943 |
| L4_*mut-16*_25C_2_ small_RNA | 12,592,284 | 10,981,746 | 2,076,206 |

**Table S2. Oligonucleotide sequences.** Related to qPCR experiments in Methods.

| **Primer name** | **Sequence** |
|---|---|
| **AR.020 *rpl-32* qPCR F** | CAAGGTCGTCAAGAAGAAGC |
| **AR.021 *rpl-32* qPCR R** | GGCTACACGACGGTATCTGT |
| **AR.030 *rab-3* qPCR F** | GCCTTCGTCTCTACTGTCGG |
| **AR.031 *rab-3* qPCR R** | CGGCGGTATCCCAGATTTGA |
| **AR.032 *myo-3* qPCR F** | GCCTACGCTGATGCTCAGAA |
| **AR.033 *myo-3* qPCR R** | CCTTCTGGCGTTGTTCCTCT |
| **AR.036 *faah-1* qPCR F** | TTCACACCAACACCTGCACT |
| **AR.037 *faah-1* qPCR R** | TGGAATGACTGTATGTCCGGC |
| **AR.050 *alg-3* qPCR F** | GGATCTGGTTCACTGTCACC |
| **AR.051 *alg-3* qPCR R** | CGACAGGAGGTGATAAAGATCC |
| **AR.052 *alg-4* qPCR F** | CCTGCCACTTGCCGTAGTTA |
| **AR.053 *alg-4* qPCR R** | CCTTCGATCTTCCGTACTTCAC |

**Table S3. Pseudopod formation counts.** Related to *in vitro* spermatid activation assay in Methods.

| | | | **Activated** | | | |
|---|---|---|---|---|---|---|
| **Sample** | **Pronase E** | **Unactivated** | **Normal** | **Short** | **Double** | **Spiky Intermediate** |
| **20ºC wild-type** | - | 200 | 0 | 0 | 0 | 0 |
| **20ºC wild-type** | + | 11 | 189 | 0 | 0 | 0 |
| **25ºC wild-type** | - | 200 | 0 | 0 | 0 | 0 |
| **25ºC wild-type** | + | 29 | 129 | 44 | 0 | 0 |
| **20ºC *mut-16*** | - | 200 | 0 | 0 | 0 | 0 |
| **20ºC *mut-16*** | + | 22 | 125 | 47 | 6 | 0 |
| **25ºC *mut-16*** | - | 200 | 0 | 0 | 0 | 0 |
| **25ºC *mut-16*** | + | 29 | 28 | 73 | 0 | 70 |
